# PDGFD-induced pericyte contractility drives pathological neoangiogenesis

**DOI:** 10.64898/2026.10.28.762436

**Authors:** Andreia Pena, Lenka Henao Misikova, Yi Jin, Marta P. Saraiva, Daniela Ramalho, Catarina Esteves, Marie Ouarné, Marta Martins, Yulia Carvalho, Erika B. Folestad, Ulf Eriksson, Lars Jakobsson, Ana M. Figueiredo, Cláudio A. Franco

## Abstract

Neovascular tufts (NVTs) are a defining feature of proliferative retinopathies such as diabetic retinopathy and retinopathy of prematurity, in which they compromise retinal integrity and vision. Excessive endothelial cell (EC) sprouting underlies their formation, yet the cellular and mechanical mechanisms that shape NVTs, and the contribution of pericytes (PCs) to this process, remain unclear. In this work, we investigated the role of actomyosin contractility in ECs and PCs during pathological angiogenesis and NVT formation. We identified non-muscle myosin IIA (NMIIA) and phosphorylated myosin light chains (MLCs) as a key signature of PCs and ECs in retinopathy and NVTs. Using inducible deletion of *Myh9* in the oxygen-induced retinopathy mouse model, we uncover distinct and opposing roles for NMIIA in the two cell types. Endothelial NMIIA promotes neovascularisation and prevents enlargement of NVTs, whilst PC NMIIA drives neovascular tuft formation. Mechanistically, we identify platelet-derived growth factor D (PDGFD), but not PDGFB, as a driver of pathological PC actomyosin contractility through Rho-associated coiled-coil kinase (ROCK). Genetic deletion of *Pdgfd* partially phenocopies the loss of NMIIA in PCs, reducing NVT formation and lowering NMIIA and phosphorylated MLC levels within NVTs. Our findings establish cell-type-specific actomyosin contractility as a determinant of NVT pathogenesis and identify PDGFD-ROCK-NMIIA signalling in PCs as a therapeutic target in proliferative retinopathies.

## INTRODUCTION

Retinopathies represent a major cause of vision loss worldwide and are highly associated with vascular dysfunction [1–3]. These conditions, including diabetic retinopathy (DR) and retinopathy of prematurity (ROP), are typically characterised by two distinct stages: an initial non-proliferative phase, dominated by vascular regression, capillary dropout, and gradual ischemia, and a subsequent proliferative stage distinguished by the pathological growth of new vessels in response to hypoxic insult [3–5]. A crucial and unique feature of proliferative retinopathies is the emergence of neovascular tufts (NVTs), which are glomeruloid-like vascular structures that protrude into the vitreous. These vessels are particularly unstable and often result in haemorrhage, oedema, and fibrosis, all of which contribute directly to retinal detachment and irreversible vision impairment [4, 5]. A major therapeutic challenge is to block NVT formation while simultaneously improving retinal revascularisation and vascular function. Current therapies for proliferative retinopathies preferentially target vascular endothelial growth factor (VEGF) signalling [6]. While these interventions have significantly improved clinical outcomes, they are not restorative and are often associated with incomplete or transient responses and potential adverse effects on physiological vessels [6]. Thus, elucidating the mechanisms underlying NVT formation and pathology may lead to more targeted and effective therapeutic approaches. Remarkably, very limited information has arisen on the cellular and molecular mechanisms of NVT formation.

Dysfunction of endothelial cells (ECs) and pericytes (PCs) contributes to vascular pathologies, including retinopathies [1, 2]. ECs exhibit aberrant proliferation, loss of polarity, and loss of junctional integrity, all of which contribute to NVT formation and vascular leakage [7–9]. PC loss and dysfunction are also hallmarks of DR and ROP, compromising barrier integrity and promoting vascular leakage [10–14]. Single-cell RNA sequencing in the oxygen-induced retinopathy (OIR) model, an established murine model of retinopathy [15], revealed a unique PC type signature associated with capillary dysfunction [16]. Signalling pathways such as PDGFB/PDGFRβ and TGF-β/ALK1/ALK5 regulate PC recruitment and stability [10, 11]. In addition, targeting the PDGFRβ/NCK1/2 axis or serum response factor (SRF) signalling can reduce NVT formation and improve vascular repair in ischemic retinopathy [12, 13].

Recently, proteomic analyses of OIR mouse retinas and proliferative DR (PDR) human samples identified several upregulated cytoskeletal proteins [17–19]. Amongst the different cytoskeletal-related proteins, there was a strong correlation between Myh9 expression, coding for the heavy chain of the non-muscle myosin II A complex (NMIIA) and vascular pathology [17–19]. Remarkably, NMIIA was highly enriched in both ECs and PCs [19].

Actomyosin-based contraction, driven by non-muscle myosin II (NMII) proteins, plays a central role in controlling cell shape, migration, adhesion, and contractility [20–22]. NMII exerts its function by cross-linking and mechanical contraction of actin filaments [23–25]. In ECs, NMIIA regulates protrusive activity and invasion during sprouting angiogenesis, primarily by controlling the formation of dactylopodia [20]. However, NMIIA function in PCs remains unexplored. Taken together, we hypothesise that NMIIA may play a pivotal role in regulating both EC and PC contractility during pathological neovascularisation.

In this study, we dissected the role of NMIIA in ECs and PCs during pathological neovascularisation, using the mouse OIR model. Remarkably, *Myh9* inactivation shows a dichotomous role in NVT formation. Whilst its deletion in ECs leads to enlarged NVTs, inactivation in PCs leads to a reduced number and size of these pathological lesions. Moreover, we identify PDGFD as a master regulator of PC contractility in NVT formation, acting through ROCK-dependent phosphorylation of myosin regulatory light chains. Together, our findings highlight an unprecedented role of NMIIA and actomyosin contractility in the regulation of retinopathies and identify NMIIA as a key regulator of EC-PC interactions during retinal neovascularisation and as a promising therapeutic target for treating ischemic retinopathies.

## RESULTS

### NMIIA is upregulated in ECs and PCs during ischemic retinopathy

The OIR model is widely used to study pathological retinal neovascularization and recapitulates several aspects of ROP and PDR [26, 27]. In this model, mice are exposed to hyperoxia at early postnatal stages, which disrupts normal retinal vascular development and causes central vaso-obliteration [28]. Upon return to normoxia, the resulting hypoxia, caused by vaso-obliteration, triggers VEGFA overexpression and pathological neoangiogenesis, including the formation of abnormal and leaky NVTs. These glomeruloid-like vascular structures peak around 5 days after return to normoxia. Proteomic profiling identified an upregulation of multiple actin cytoskeleton-related proteins in OIR mouse retinas and PDR patients [17–19]. From the different actin-related proteins, NMIIA was reported to be highly expressed in ECs and PCs in pathological vessels, including NVTs [19], suggesting that alterations in vascular actomyosin contractility may contribute to disease onset and/or progression.

First, we took an unbiased approach by using publicly available single-cell RNA sequencing (scRNAseq) datasets of OIR mouse retinas (GSE150703 and GSE216676; [29, 30]) to evaluate the expression levels of all myosin heavy and light chains (Fig.S1). ECs and PCs were identified based on the expression of the canonical markers Cldn5, Pecam1, and Cdh5 for ECs, and Pdgfrb, Rgs5, Kcnj8, and Vtn for PCs [31]. At this resolution, ECs and PCs were combined in a single cluster and showed that this cluster has a high level of expression for heavy and light chains (Fig.S1B). We subsequently integrated the two datasets and annotated the cells as ECs, PCs, as well as proliferating ECs or proliferating PCs, using Mki67 and Top2a as proliferation markers (Fig.1A-B, Fig.S2). Finally, we compared the average expression of all known myosin heavy and light chain genes in ECs and PCs between normoxic and OIR conditions (Fig. 1C). Myh9 was the myosin heavy chain with the highest expression amongst all myosin heavy chains in both ECs and PCs and was upregulated in OIR conditions (Fig.1C). ECs express high levels of myosin light chains (Myls) Myl6, Myl12a and Myl12b, whilst PCs express high levels of Myl6, Myl12a and Myl9. Amongst these, only Myl6 was clearly increased in OIR in both ECs and PCs, whereas Myl9, Myl12a and Myl12b were highly expressed but not consistently induced (Fig.1C).

**Figure 1.**
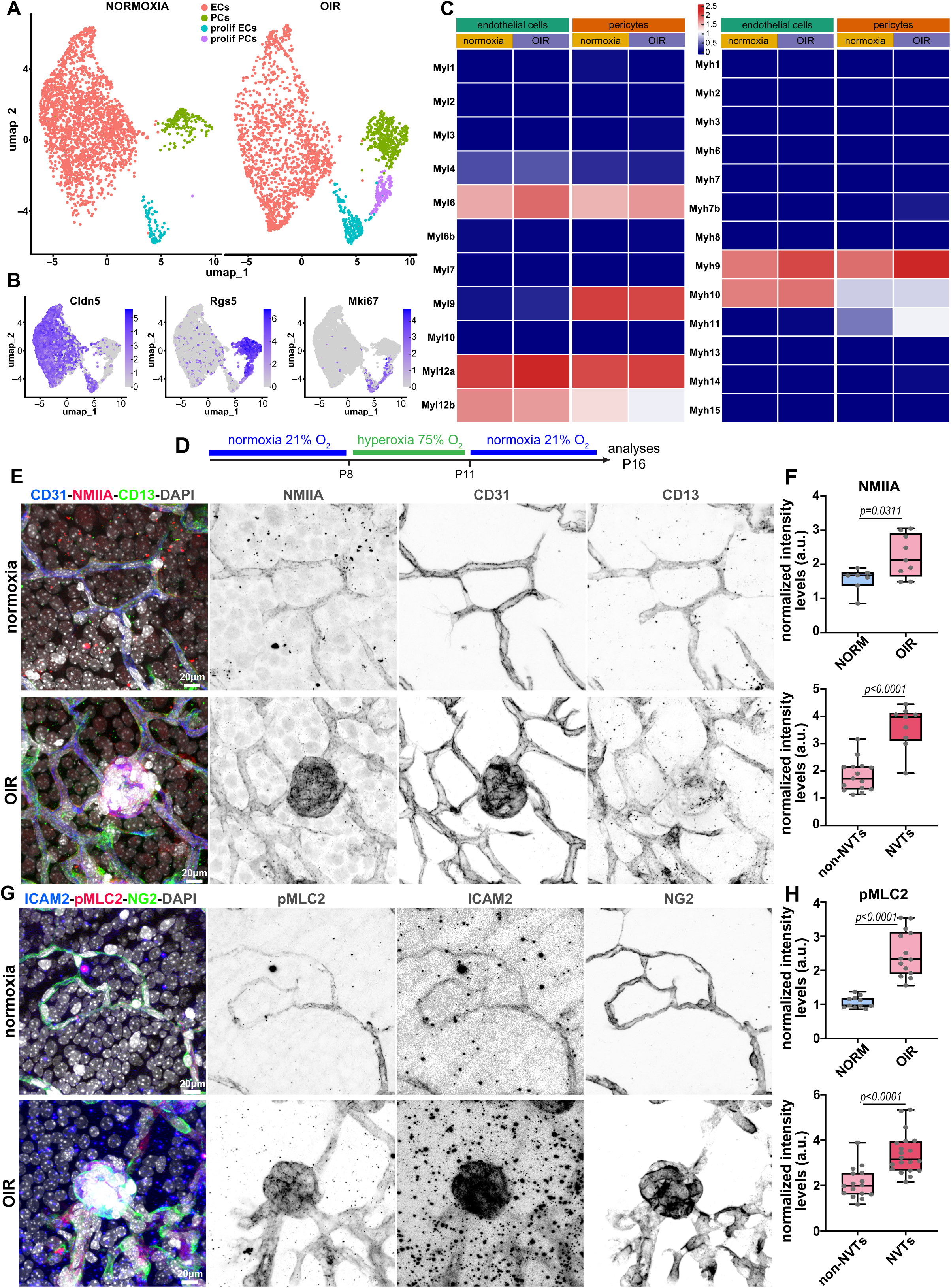
NMIIA and its activity are upregulated in ECs and PCs during OIR, in particular in NVTs. **A)** Uniform manifold approximation and projection (UMAP) of integrated ECs and PCs subclusters from normoxic and oxygen-induced retinopathy (OIR) retinas during neovascularisation (GEO accession numbers GSE150703 and GSE216676) annotated as ECs, PCs, proliferating ECs (prolif ECs) and proliferating PCs (prolif PCs). **B)** UMAP plots showing expression of canonical markers Cldn5 (ECs), Rgs5 (PCs), and Mki67 (proliferative cells) across integrated clusters. **C)** Heatmaps of average expression levels of myosin light chain (Myl; left) and heavy chain (Myh; right) genes in ECs and PCs under normoxic and OIR conditions. **D)** OIR timeline scheme. **E)** Representative images of retinal capillaries under normoxic or OIR conditions from wild-type mice at P16, stained for ECs (CD31, blue), PCs (CD13, green), NMIIA (red) and nuclei (DAPI, grey). Single channels are shown as inverted greyscale. **F)** Quantification of NMIIA levels, normalised to vessel area, in normoxic (NORM) versus OIR retinas (top; each dot represents one image; N=7 NORM; N=9 OIR), and in non-NVT vessels versus NVT vessels of OIR retinas (bottom; each dot represents one vessel segment or NVT; N=15 non-NVT; N=9 NVT). **G)** Representative images of retinal capillaries and of a NVT under normoxic or OIR conditions from wild-type mice at P16, stained for ECs (ICAM2, blue), pMLC2 (red), PCs (NG2, green) and nuclei (DAPI, grey). Single channels are shown as inverted greyscale. **H)** Quantification of pMLC2 levels, normalised to vessel area, in normoxic (NORM) versus OIR retinas (top; each dot represents one image; N=11 NORM; N=14 OIR), and in non-NVT vessels versus NVT vessels of OIR retinas (bottom; each dot represents one vessel segment or NVT; N=16 non-NVT; N=20 NVT). Data are presented as box plots with whiskers, min to max and median, from at least 3 independent litters.

Because the activity of the regulatory light chains is primarily controlled by phosphorylation rather than by transcription, we next evaluated both NMIIA protein levels and the phosphorylation status of the regulatory light chains in situ. First, we assessed NMIIA and pan-phosphorylated myosin light chain (pMLC2) by immunofluorescence on whole-mount retinas of wild-type animals, at the peak of NVT formation and at the corresponding time point in normoxic controls (Fig.1D). NMIIA was detected in both CD31+ ECs and CD13+ PCs, and its levels were significantly higher in OIR than in normoxia (Fig.1E,F). Importantly, within OIR retinas, NMIIA levels were markedly higher in NVTs than in the adjacent non-tuft vasculature (Fig.1E,F). Similarly, pMLC2 levels were significantly increased in OIR compared to normoxia and were strongly enriched in NVTs relative to non-tuft vessels (Fig.1G,H). To further validate these findings at the single cell-level, we crossed *Cdh5-Cre^ERT2^* (EC-specific) and *Pdgfrβ-Cre^ERT2^* (MC-specific) mouse lines with the *Rosa26^mTmG^* reporter line. Mice were injected with a low dose of 4-hydroxytamoxifen (TAM) 3 days before sacrifice to sparsely label single ECs or MCs, respectively, and NMIIA expression was assessed at the peak of NVT formation and at the corresponding time point in normoxic controls (Fig.S3). Consistent with scRNAseq data, NMIIA levels were significantly upregulated in ECs and PCs in OIR conditions compared to normoxia (Fig.S3B-E). Interestingly, the high levels of NMIIA and activated MLC coincided with a marked change in the morphology of NVT-associated ECs and PCs. Instead of a more spindle-shaped or slightly branched morphology seen in non-NVT vessels (OIR or non-OIR retinas), NVT-associated ECs and PCs showed a more rounded shape and exhibited blebs, reminiscent of highly contractile cells (Fig.S3B and D, and Supplementary Videos 1-7). Together, these results highlight a possible crucial role for NMIIA and actomyosin contractility in pathological angiogenesis in ECs and/or PCs.

### EC NMIIA promotes neovascularization and limits NVT formation

Next, we sought to test the role of endothelial NMIIA for OIR-induced neoangiogenesis. We and others previously demonstrated that NMIIA regulates EC dactylopodia formation and invasive behaviour during physiological angiogenesis [20, 32]. NMIIA EC-KO retinas showed decreased radial expansion, increased vessel density, and increased tip cell numbers [20]. To address the role of EC NMIIA in OIR-associated neoangiogenesis, we crossed *Myh9*^flox^ mice with the *Cdh5-Cre^ERT2^* strain. To avoid confounding effects of earlier defects in angiogenesis, we induced endothelial-specific deletion of *Myh9* (*Myh9*^EC-iKO^) immediately after returning to normoxia, via TAM injections at P11 and P13 and pups were sacrificed at P16 (Fig.2A). Loss of NMIIA protein in OIR mice was confirmed by a significant reduction in NMIIA immunofluorescence intensity in ECs of *Myh9*^EC-iKO^ retinas compared to controls (Fig.S4A,B), as reported previously [20]. At P13, immediately after return to normoxia, the extent of hyperoxia-induced vaso-obliteration was indistinguishable between *Myh9^EC-WT^* and *Myh9^EC-iKO^* retinas (Fig.S4C,D). However, by P16, *Myh9^EC-iKO^* retinas exhibited a significantly higher avascular area when compared to Cre-negative wild-type littermates (*Myh9*^EC-WT^), suggesting impaired retinal revascularisation following endothelial Myh9 deletion (Fig. 2B–D). This is consistent with the relevant role of NMIIA for endothelial tip cell invasion [20, 21]. However, the area and size of NVTs were significantly increased in *Myh9*^EC-iKO^ retinas relative to controls (Fig.2C-F). These defects were not associated with a loss of MCs. NG2 coverage of NVTs was unchanged (Fig.2G), desmin coverage of NVTs was slightly increased, and both NG2 and desmin coverage of neo-angiogenic sprouts were unaffected (Fig.S4E,F). Together, these results support that NMIIA is a key regulator of EC invasive behaviour, both in physiological and pathological angiogenesis. We previously demonstrated that NMIIA loss in ECs decreases EC proliferation [20], which could contribute to decreased neovascularisation, but would have an opposite effect on NVT enlargement. Taken together, we postulate that EC NMIIA function is important for efficient neovascularisation and limits the formation and expansion of NVTs.

**Figure 2.**
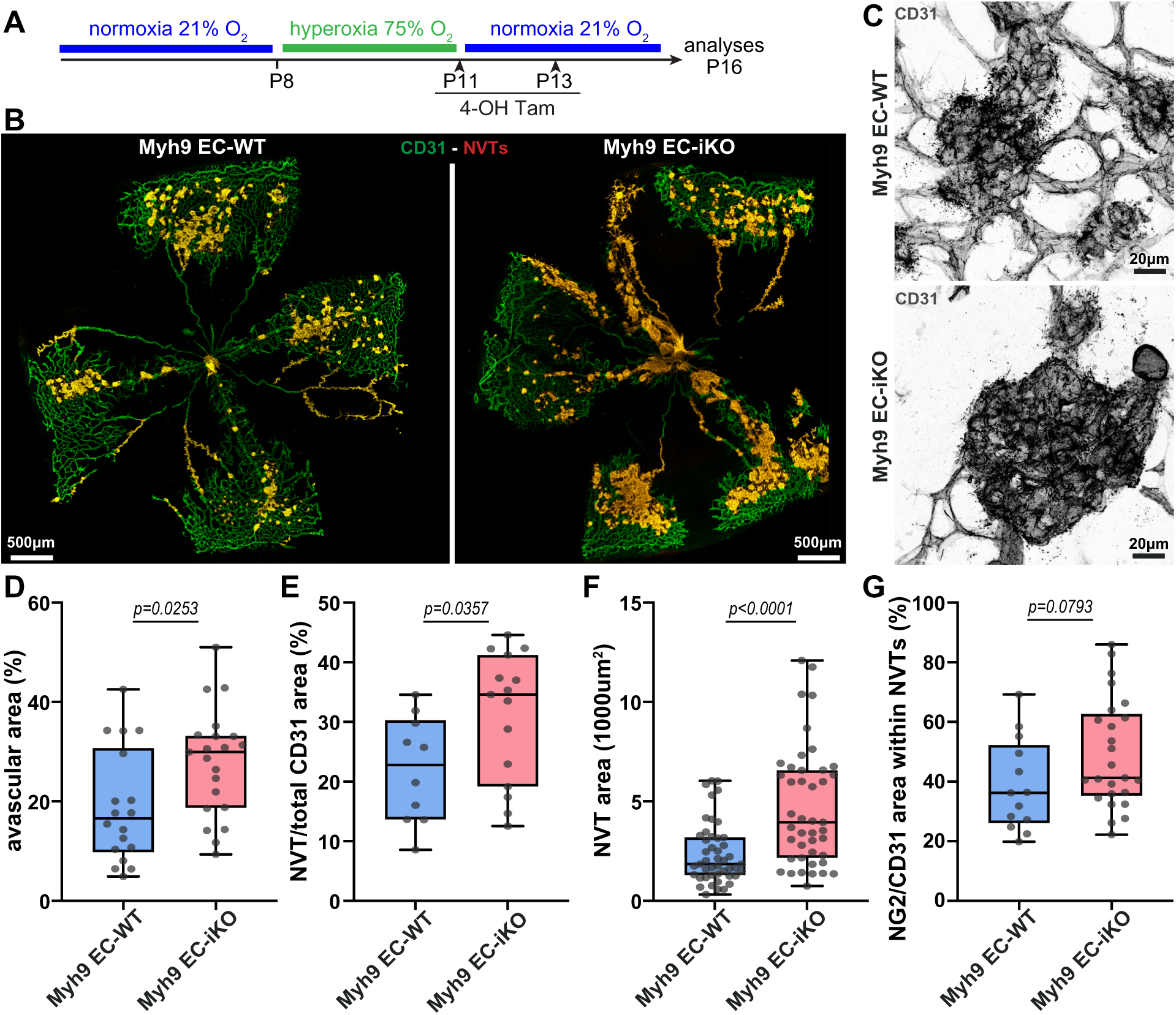
Endothelial *Myh9* deletion increases pathological neovascularization in OIR. **A)** Experimental timeline for EC-specific *Myh9* deletion during OIR. **B)** Representative images of P16 OIR retinas from *Myh9*^EC-WT^ and *Myh9*^EC-iKO^ mice stained for CD31 (green). Neovascular tufts (NVTs) were manually delineated (red) and superimposed to CD31 staining. **C)** Higher-magnification images of NVTs stained for CD31 (grey) from *Myh9*^EC-WT^ and *Myh9*^EC-iKO^ mice. **D)** Quantification of the percentage of avascular area in P16 OIR retinas from *Myh9*^EC-WT^ and *Myh9*^EC-iKO^ mice. Each dot represents one mouse retina. N=18 *Myh9*^EC-WT^; N=21 *Myh9*^EC-iKO^. **E)** Quantification of the total NVT burden per retina, calculated as the sum of all NVT areas divided by the whole-retina CD31+ area (NVT/CD31 areas), from *Myh9*^EC-WT^ and *Myh9*^EC-iKO^ mice. Each dot represents one mouse retina. N=10 *Myh9*^EC-WT^; N=15 *Myh9*^EC-iKO^. **F)** Quantification of individual NVT area in P16 OIR retinas from *Myh9*^EC-WT^ and *Myh9*^EC-iKO^ mice. Each dot represents one NVT. N=46 *Myh9*^EC-WT^; N=43 *Myh9*^EC-iKO^. **G)** Quantification of the percentage of PC coverage (NG2+) in individual NVTs (CD31+) of P16 OIR retinas from *Myh9*^EC-WT^ and *Myh9*^EC-iKO^ mice. Each dot represents one NVT. N=12 *Myh9*^EC-WT^; N=25 *Myh9*^EC-iKO^. All data are presented as box plots with whiskers, min to max and median, from at least 3 independent litters.

### PC NMIIA regulates developmental angiogenesis

In contrast to endothelial NMIIA, the role of PC actomyosin contraction in blood vessel development has not yet been assessed. Thus, first, we analysed the role of PC NMIIA in normal retinal development. To do so, we crossed the *Myh9*^flox^ mouse line with the Pdgfrβ-Cre^ERT2^ line, which allows for Cre recombinase-mediated gene deletion in MCs (PCs and vSMCs). The MC-specific deletion of *Myh9* was performed by intraperitoneal injections of TAM at P1, P2 and P3, and we focused our analysis on the retinal vasculature at P6 (Fig.3A). As controls, we used the Cre-negative littermates (*Myh9*^MC-WT^). Two distinct regions can be identified at P6: i) the sprouting front (SF) region, where ECs proliferate extensively, expand the network generating immature vessels and recruit new PCs; and ii) the remodelling plexus (RP) region, closer to the optic nerve, where vessel remodelling is ongoing and a hierarchical network with arteries, arterioles, venules and veins starts emerging [33] (Fig.S5).

**Figure 3.**
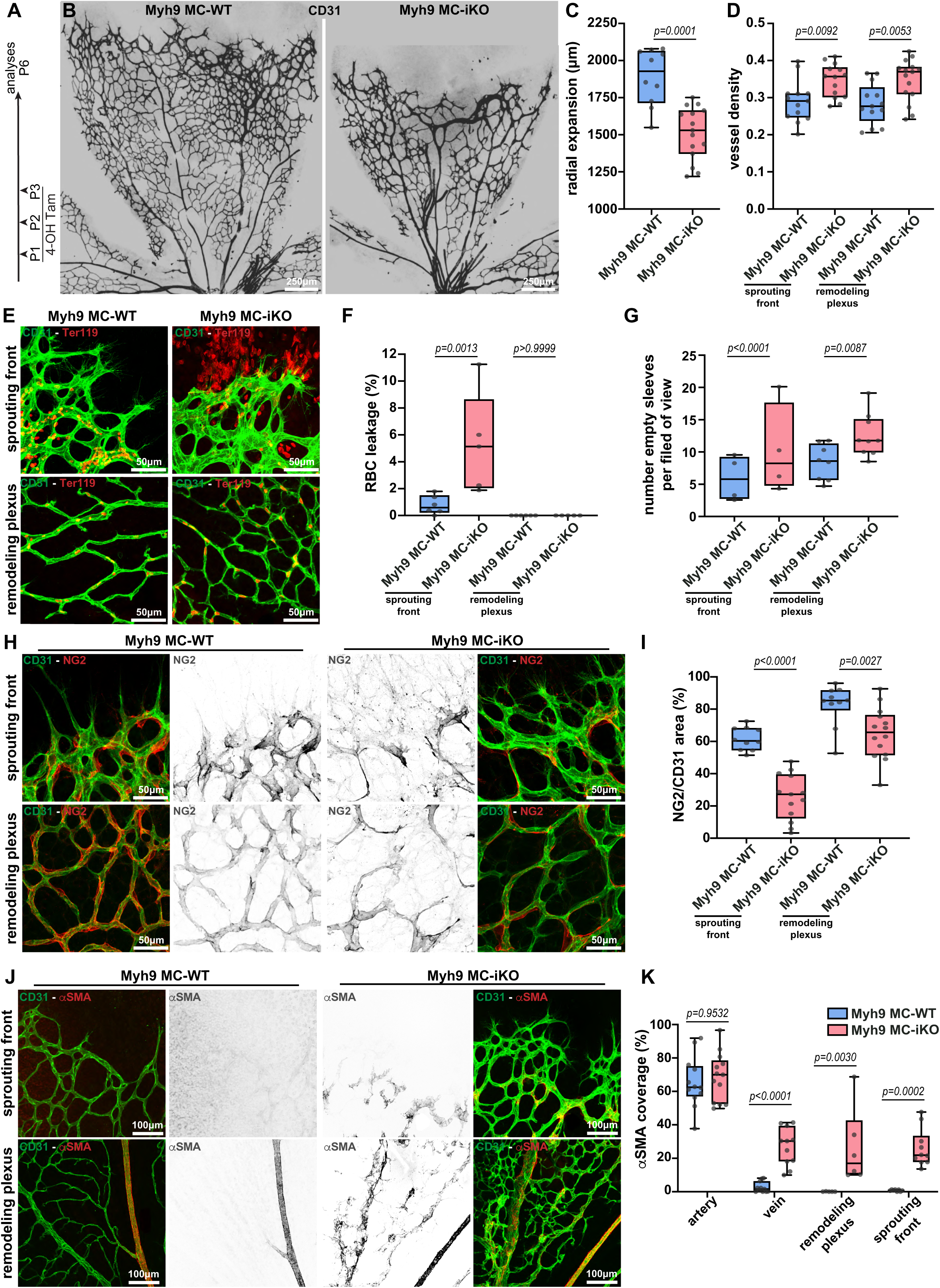
PC NMIIA is required for angiogenesis and vascular stability. **A)** Experimental timeline for MC-specific *Myh9* deletion under physiological postnatal stages. **B)** Representative images of blood vessels (CD31, grey) of P6 retinas from *Myh9*^MC-WT^ and *Myh9*^MC-iKO^ mice. **C)** Quantification of radial expansion in P6 retinas from *Myh9*^MC-WT^ and *Myh9*^MC-iKO^ mice. Each dot represents one retina. N=10 *Myh9*^MC-WT^; N=15 *Myh9*^MC-iKO^. **D)** Quantification of vessel density at SF or RP in P6 retinas from *Myh9*^MC-WT^ and *Myh9*^MC-iKO^ mice. Each dot represents one retina. N=13 *Myh9*^MC-WT^; N=14 *Myh9*^MC-iKO^. **E)** Representative images of SF and RP of P6 retinas from *Myh9*^MC-WT^ and *Myh9*^MC-iKO^ retinas. Retinas were stained for vessels (CD31, green) and red blood cells (RBCs, Ter-119, red). **F)** Quantification of RBC leakage at the SF and RP of P6 retinas from *Myh9*^MC-WT^ and *Myh9*^MC-iKO^ mice. Each dot represents one retina. N=6 *Myh9*^MC-WT^; N=5 *Myh9*^MC-iKO^. **G)** Quantification of collagen empty sleeves at the SF and RP of P6 retinas from *Myh9*^MC-WT^ and *Myh9*^MC-iKO^ mice. Each dot represents one retina. SF, N=4 for both *Myh9*^MC-WT^ and *Myh9*^MC-iKO^. RP, N=8 for *Myh9*^MC-WT^; N=9 *Myh9*^MC-iKO^. **H)** Representative images of SF and RP of P6 retinas from *Myh9*^MC-WT^ and *Myh9*^MC-iKO^ retinas stained for vessels (CD31, green) and PCs (NG2, red and grey). **I)** Quantification of PC coverage (NG2+) in vessels (CD31+) in the SF or RP of P6 retinas from *Myh9*^MC-WT^ and *Myh9*^MC-iKO^ mice. Each dot represents one retina. N=10 *Myh9*^MC-WT^; N=13 *Myh9*^MC-iKO^. **J)** Representative images of SF and RP of P6 retinas from *Myh9*^MC-WT^ and *Myh9*^MC-iKO^ stained for vessels (CD31, green) and αSMA (red and grey). **K)** Quantification of αSMA-coverage in vessels (CD31+) in arteries, veins, in the SF or RP of P6 retinas from *Myh9*^MC-WT^ and *Myh9*^MC-iKO^ mice. Each dot represents one vessel/capillary region. Artery: N=12 *Myh9*^MC-WT^, N=13 *Myh9*^MC-iKO^; Vein: N=10 *Myh9*^MC-WT^, N=11 *Myh9*^MC-iKO^; RP: N=6 *Myh9*^MC-WT^, N=6 *Myh9*^MC-iKO^; SF: N=7 *Myh9*^MC-WT^, N=10 *Myh9*^MC-iKO^. All data are presented as box plots with whiskers, min to max, with median from at least 3 independent litters.

To assess recombination efficiency, we intercrossed the *Rosa26^mTmG^* reporter line [34] with the *Myh9^flox^::Pdgfrβ-Cre^ERT2^* line. In this setup, Cre-positive Myh9^wt/wt^ (*Myh9*^wt/wt^::*Rosa26^mTmG^*::Pdgfrβ-Cre^ERT2^) animals served as controls. Quantitative analysis of single recombined MCs showed a significant reduction in NMIIA levels at the SF and in the RP in *Myh9*^MC-iKO^ (Fig.S5A,B). Interestingly, this analysis also revealed that PCs at the sprouting front express significantly higher levels of NMIIA than PCs in the RP of control P6 mouse retinas (Fig.S5B), suggesting that SF PCs may require a more robust actomyosin cytoskeleton.

Next, we investigated the vascular consequences of *Myh9* deletion in MCs. Remarkably, *Myh9*^MC-iKO^ showed an aberrant vasculature at P6 (Fig.3B and Fig.S5C). The phenotype was characterised by reduced radial outgrowth (Fig.3C) and increased vessel density in both the SF and the RP (Fig.3D and Fig.S5C). Contrary to the phenotype of deleting *Myh9* in ECs [20], no significant differences were observed in the number of endothelial tip cells or their filopodia number between *Myh9*^MC-WT^ and *Myh9*^MC-iKO^ retinas (Fig.S5D). In the SF region, we observed a large dilation of blood vessels (Fig.S5E). Vein diameter was also increased, whereas the diameter of arteries and of RP capillaries was unchanged (Fig.S5E). This phenotype was associated with leaky, unstable vessels, highlighted by an increase in the number of collagen IV empty sleeves, a surrogate for excessive vascular pruning [35, 36], and an increase in extravascular red blood cells (micro-haemorrhages) (Fig.3E-G and Fig.S5F). However, no major differences were apparent in EC junction structure (Fig.S5G). These phenotypes are reminiscent of decreased PC coverage of blood vessels, which leads to EC hyperplasia and increased permeability [37, 38].

Thus, we investigated the coverage of retinal vessels by PCs and vSMCs using canonical MC markers, NG2, desmin, PDGFRβ, and αSMA. In *Myh9*^MC-iKO^ retinas, the three PC markers NG2, desmin and PDGFRβ were significantly reduced at the SF (Fig.3H,I and Fig.S6A,B). Similarly, within the RP, NG2 and PDGFRβ coverage were decreased, whereas desmin coverage remained unchanged (Fig.3H,I and Fig.S6A,B). Remarkably, αSMA, a well-established vSMC marker, showed a significant increase in veins and in capillaries of both the SF and the RP (Fig.3J,K). Under physiological conditions, quiescent PCs do not express αSMA. However, PCs can become activated by biomechanical and inflammatory stimuli [10]. This was already reported in several disease mouse models, including ischemic retinopathy [12], diabetic retinopathy [39], and pancreatic ductal adenocarcinoma [40], and lineage tracing in OIR demonstrated that αSMA+ PCs are derived from *bona fide* PCs [12].To test whether this phenotype was transient, we collected retinas at P12 and observed that it became stronger over time, characterised by severely abnormal vessel morphology, particularly at the SF, including arteriovenous shunt-like vessels (Fig. S6C) decorated with αSMA-positive MCs (Fig. S6D). These malformations are reminiscent of defects reported for SRF deletion in PCs and PDGFB deletion in ECs [13, 41].

Altogether, our findings indicate that NMIIA in PCs is essential for blood vessel development and vessel stability during sprouting angiogenesis. We concluded that NMIIA is required for the adequate coverage of nascent blood vessels by PCs, in particular at the sprouting front.

### NMIIA is essential for PC migration

Next, we aimed to understand the underlying cellular mechanism of decreased PC coverage, particularly in the SF region. Firstly, we investigated PC and EC apoptosis by analysing cleaved caspase-3 staining in *Myh9*^MC-iKO^ and control mice. No differences were detected (Fig.S7A,B). Secondly, we assessed the number of proliferative PCs and ECs. For this analysis, PCs were identified as NG2-positive cells and ECs were identified by their ERG-positive nuclei, and EdU incorporation was scored separately in each population (Fig.4A,B and Fig.S7C). In *Myh9*^MC-iKO^ retinas, we observed a notable reduction in the proliferation of PCs at the SF, but no significant differences in RP when compared to their control littermates (Fig.4A,B). A trend of reduced EC proliferation, albeit not statistically significant, was also observed at the SF of *Myh9*^MC-iKO^ retinas (Fig.4B and Fig.S7C), yet this did not translate into changes in EC density at either the SF or RP (Fig.S7D). This suggests that the absence of *Myh9* has a selective impact on PC proliferation during angiogenesis, primarily affecting the SF.

**Figure 4.**
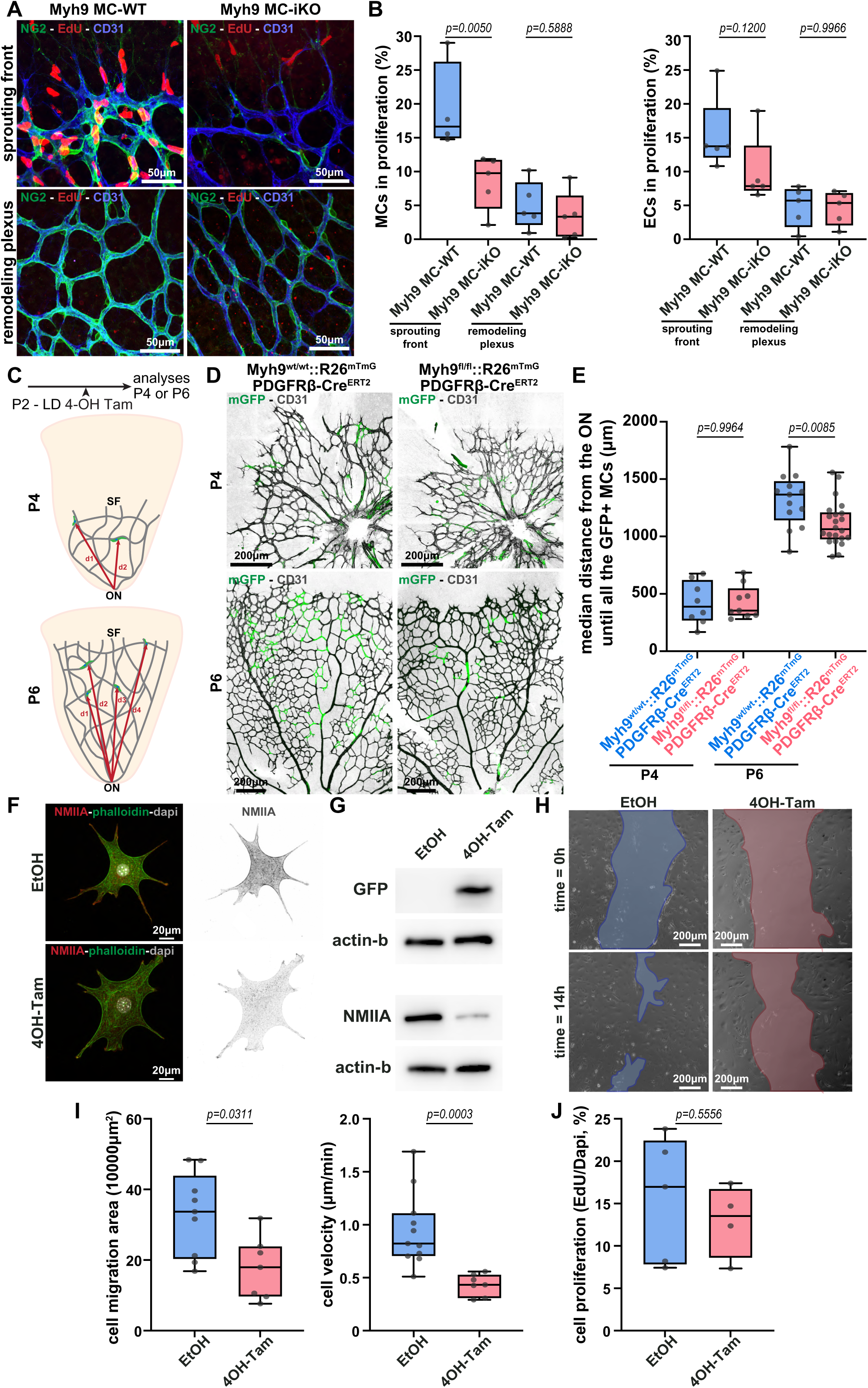
NMIIA regulates MC migration and proliferation in mouse retinas. **A)** Representative images of SF and RP of P6 retinas from *Myh9*^MC-WT^ and *Myh9*^MC-iKO^ mice stained for vessels (CD31, blue), PCs (NG2, green), and proliferating cells (EdU, red). **B)** Quantification of proliferating MCs (left panel) and ECs (right panel) in the SF and RP of P6 retinas from *Myh9*^MC-WT^ and *Myh9*^MC-iKO^ mice. Each dot represents one mouse retina. N=5 mice for both *Myh9*^MC-WT^ and *Myh9*^MC-iKO^. **C)** Experimental timeline for *in vivo* migration assessment. SF: sprouting front; ON: optic nerve. **D)** Representative images of P4 (upper panels) or P6 (bottom panels) retinas from *Myh9^wt/wt^::Pdgfrβ-Cre^ERT2^::R26^mTmG^* and *Myh9^fl/fl^::Pdgfrβ-Cre^ERT2^::R26^mTmG^* mice stained for vessels (CD31, grey) and mGFP (green, recombined PCs). **E)** Quantification of the median distance of mGFP+ PCs from the ON in P4 or P6 retinas from *Myh9^wt/wt^::Pdgfrβ-Cre^ERT2^::R26^mTmG^* and *Myh9^fl/fl^::Pdgfrβ-Cre^ERT2^::R26^mTmG^* mice. Each dot represents one mouse retina. P4: N=8 *Myh9^wt/wt^::Pdgfrβ-Cre^ERT2^::R26^mTmG^*, N=9 *Myh9^fl/fl^::Pdgfrβ-Cre^ERT2^::R26^mTmG^*; P6: N=13 *Myh9^wt/wt^::Pdgfrβ-Cre^ERT2^::R26^mTmG^*, N=22 *Myh9^fl/fl^::Pdgfrβ-Cre^ERT2^::R26^mTmG^* mice. **F)** Representative images of single mouse brain MCs treated with ETOH or 4OH-Tam. Cells were stained for actin (phalloidin, green), Myh9 (NMIIA, red), and nuclei (DAPI, grey). **G)** Images of western blot showing protein levels of NMIIA, GFP, and β-actin in mouse brain MCs treated with EtOH or 4OH-Tam. **H)** Representative images from a scratch-wound assay performed in mouse brain MCs treated with ETOH (left) or 4OH-Tam (right) at 0 and 14 hours. The wound area is shown at both time points. **I)** Quantification of cell migration area (left) and cell velocity (right) of mouse brain MCs treated with ETOH or 4OH-Tam. Each dot represents one well. N=10 for EtOH, N=6 for 4OH-Tam. **J)** Quantification of proliferating mouse brain MCs treated with ETOH or 4OH-Tam. Each dot represents one coverslip. N=5 for EtOH, N=4 for 4OH-Tam. All data are presented as box plots with whiskers, min to max with median, from at least 3 independent litters.

In parallel, we investigated whether impaired PC migration could also contribute to reduced PC coverage at the SF. *In vivo*, PC migration is guided by a gradient of PDGFB produced by ECs, in particular, endothelial tip cells at the SF [42]. To assess whether cell migration alone, or also cell polarity, was affected, we used the Rosa26^GNrep^ mouse reporter line [43], which allows for the identification of front-rear polarity of cells by labelling the nucleus and Golgi apparatus [33, 35]. To do so, we crossed the *Myh9*^MC-iKO^ line with Rosa26^GNrep^ mice to obtain *Myh9*^wt/wt^::Rosa26^GNrep^::Pdgfrβ-Cre^ERT2^ (controls) and *Myh9*^fl/fl^::Rosa26^GNrep^::Pdgfrβ-Cre^ERT2^ (KO) pups. We injected pups with a full dose of TAM for 3 consecutive days, starting at P1 and collected the retinas at P6 (Fig.S8A). Thereafter, we measured polarity of PCs using the same methodologies used for EC polarity analysis established previously [44–46]. Remarkably, PCs in control retinas robustly polarise towards the angiogenic front at the SF but also in the RP, suggesting that cues from the angiogenic front may have a long-range effect in the mouse retina (Fig.S8B,C). Yet, *Myh9*^fl/fl^::Rosa26^GNrep^::Pdgfrβ-Cre^ERT2^ retinas showed no differences in the polarity pattern of PCs, at the SF or RP, when compared to control retinas (Fig.S8B,C). This suggests that NMIIA-deficient PCs retain the ability to read PDGFB gradients. Next, we performed an analysis of the distribution of genetically-labelled PCs over time, as a proxy for cell migration *in vivo*. To do so, we performed pulsed-labelling of PCs using a single injection of low-dose TAM at P2, followed by analysis of Myh9^wt/wt^::Rosa26^mTmG^::Pdgfrβ-Cre^ERT2^ and Myh9^fl/fl^::Rosa26^mTmG^::Pdgfrβ-Cre^ERT2^ at P4 or P6. To assess PC migration in fixed mouse retinas, we measured the distance between the optic nerve and each GFP-positive MC. At P4, GFP-positive cells show the same distribution in the mouse retina for both conditions (Fig.4C-E). Yet, at P6, labelled MCs from *Myh9*^MC-WT^ migrated further away from the optic nerve, reaching the SF or its vicinity. Given their location, these migratory labelled cells are likely PCs. In contrast, *Myh9*^MC-iKO^ cells remained closer to the optic nerve and rarely reached the SF (Fig.4C-E).

To further confirm that NMIIA affects MC migration, we isolated primary mouse brain mural cells (mBMCs) from *Myh9^flox^::Rosa26-mTmG::Pdgfrβ-CreERT2* adult mice. These cells were then treated with either ethanol (EtOH, control) or TAM to induce deletion of Myh9, which was confirmed by immunofluorescence and western blot (Fig.4F,G). To assess MC migration, we performed a scratch-wound assay on a confluent monolayer of mBMCs previously treated with TAM or vehicle. Live imaging was carried out for 14 hours, during which the wound closure area and MC migration velocity were quantified. Our observations showed that NMIIA inactivation led to impaired MC wound closure and a decrease in MC migration speed when compared to controls (Fig.4H,I). Surprisingly, no differences were detected in MC proliferation between EtOH- and TAM-treated cells *in vitro* (Fig.4J).

Overall, these results suggest that NMIIA is essential for PC migration and proliferation, but not for their front-rear polarisation. Our data also suggest that *in vivo* proliferation defects may be a secondary effect of impaired migration and the consequent reduced exposure of PCs to EC-derived mitogenic cues at the sprouting front. Together, the defective migration and reduced proliferation lead to a marked decrease in PC coverage during angiogenesis, promoting vascular instability and vessel malformation.

### PC NMIIA is dispensable for revascularisation but promotes NVT formation

Having characterised the role of NMIIA in developmental angiogenesis, next, we sought to test the role of PC NMIIA for OIR-induced neoangiogenesis. To avoid severe developmental vascular defects arising from early loss of function of NMIIA in PCs, we induced PC-specific deletion of *Myh9* immediately after return to normoxia, via TAM injections at P11-13 (Fig.5A). Subsequently, we studied the impact of specific *Myh9* deletion on neovascular development at P16, corresponding to 5 days after return to normoxia (Fig.5A). To confirm NMIIA depletion, we performed RT-qPCR on bulk retinal RNA extraction, which revealed a significant reduction in *Myh9* mRNA levels in *Myh9*^MC-iKO^ retinas (Fig.S9A). This was accompanied by a significant decrease in NMIIA immunofluorescence signal intensity in *Myh9*^MC-iKO^ PCs compared to control PCs (Fig.S9B). Similar to the EC *Myh9* deletion, the extent of hyperoxia-induced vaso-obliteration, measured immediately after return to normoxia (P13), was indistinguishable between Myh9^MC-WT^ and Myh9^MC-iKO^ retinas (Fig.S9C,D). Next, we characterised key parameters of OIR-induced neoangiogenesis. Deletion of *Myh9* in MCs did not affect revascularisation compared with Cre-negative wild-type littermates (*Myh9*^MC-WT^) (Fig.5B-D). However, deletion of *Myh9* in MCs decreased NG2- and desmin-positive PC coverage in sprouting neovessels of *Myh9*^MC-iKO^ retinas compared to control *Myh9*^MC-WT^ retinas (Fig.S9E,F). This is consistent with the relevant role of NMIIA for PC migration in developmental angiogenesis (Fig.4). Yet, surprisingly, we also observed a significant reduction in the number and the size of NVTs in *Myh9*^MC-iKO^ in comparison to *Myh9*^MC-WT^ animals (Fig.5C,E,F). Moreover, this decrease in NVT formation was not associated with a loss of PC coverage. Analysis of PC coverage by immunofluorescence indicated similar NG2- and desmin-positive coverage of the residual NVTs in *Myh9*^MC-iKO^ and *Myh9*^MC-WT^ animals (Fig.5G and Fig.S9E,F). Taken together, these findings show that PCs require NMIIA activity to properly migrate in developmental and pathological angiogenesis, and that NMIIA-dependent PC function, rather than PC presence per se, is required for NVT formation and development.

**Figure 5.**
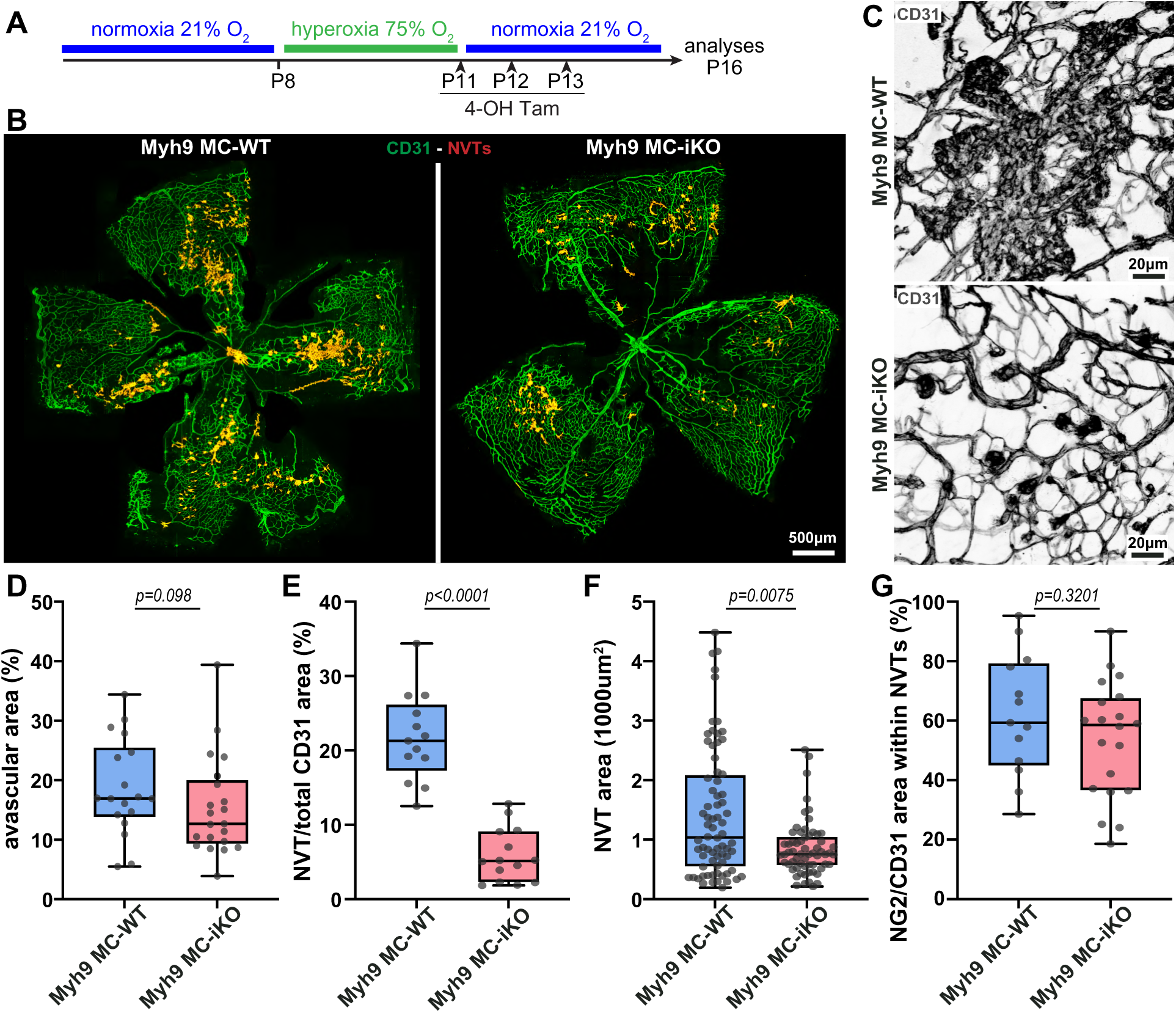
MC-specific deletion of Myh9 reduces NVT formation. **A)** Experimental timeline for MC-specific deletion of Myh9 in OIR. **B)** Representative images of P16 OIR retinas from *Myh9*^MC-WT^ and *Myh9*^MC-iKO^ mice stained for CD31 (green). Neovascular tufts (NVTs) were manually delineated (red) and superimposed on CD31 staining. **C)** Higher-magnification images of NVTs stained for CD31 (grey) from *Myh9*^MC-WT^ and *Myh9*^MC-iKO^ mice. **D)** Quantification of the percentage of avascular area in P16 OIR retinas from *Myh9*^MC-WT^ and *Myh9*^MC-iKO^ mice. Each dot represents one mouse retina. N=8 *Myh9*^MC-WT^; N=10 *Myh9*^MC-iKO^. **E)** Quantification of the total NVT burden per retina, calculated as the sum of all NVT areas divided by the whole-retina CD31+ area (NVT/CD31 areas), in P16 OIR retinas from *Myh9*^MC-WT^ and *Myh9*^MC-iKO^ mice. Each dot represents one mouse retina. N=13 *Myh9*^MC-WT^; N=14 *Myh9*^MC-iKO^. **F)** Quantification of individual NVT area in P16 OIR retinas from *Myh9*^MC-WT^ and *Myh9*^MC-iKO^ mice. Each dot represents one NVT. N=72 *Myh9*^MC-WT^; N=62 *Myh9*^MC-iKO^. **G)** Quantification of the percentage of PC coverage (NG2+) in individual NVTs (CD31+) of P16 OIR retinas from *Myh9*^MC-WT^ and *Myh9*^MC-iKO^ mice. Each dot represents one NVT. N=13 *Myh9*^MC-WT^; N=20 *Myh9*^MC-iKO^. All data are presented as box plots with whiskers, min to max, with median from at least 3 independent litters.

### PDGFD, but not PDGFB, activates PC actomyosin contractility through ROCK

Next, we sought to investigate the upstream signalling regulating actomyosin contractility in PCs during retinopathy. PDGFB/PDGFRβ signalling has been previously linked to PC migration through the activation of NCK1/2 [12]. Moreover, PDGFB is commonly identified as upregulated in the vitreous of PDR patients [47, 48]. Thus, we first tested if PDGFB/PDGFRβ signalling could activate phosphorylation of MYL9 (pMYL9), a main NMII regulatory light chain highly enriched in PCs (Fig.1C). Whilst PDGFB (40ng/mL) stimulation of hBPCs strongly activated AKT signalling, we detected a decrease in pMYL9 over time (Fig.6A), suggesting that PDGFB is not a main activator of actomyosin contractility in PCs. Yet, PDGFRβ can be activated by other PDGF family members. In a recent report, PDGFD was also identified as strongly upregulated in the vitreous of PDR patients [49], which prompted us to also test it in our *in vitro* conditions. PDGFD (40ng/mL) activated AKT signalling, although to a lesser extent than PDGFB (Fig.6A,B). In marked contrast to PDGFB, however, PDGFD induced a significant and sustained increase in the pMYL9/total MYL9 ratio (Fig.6A,B). The same conclusion was reached by immunofluorescence: PDGFD, but not PDGFB, significantly increased pMYL9 levels in hBPCs, without changing total NMIIA levels (Fig.6C,D). ROCK signalling is a central regulator of actomyosin contractility, acting upstream of myosin II activation through phosphorylation of myosin light chain (MLC) [49]. To test if ROCK activity is necessary for PDGFD-mediated activation of actomyosin contractility, we then stimulated hBPCs with PDGFD in the presence of the ROCK inhibitor Y-27632. ROCK inhibition completely abolished the PDGFD-induced increase in pMYL9 (Fig.6E,F). Our results suggest that a PDGFD-ROCK-pMLC pathway acts upstream of actomyosin contractility in PCs.

**Figure 6.**
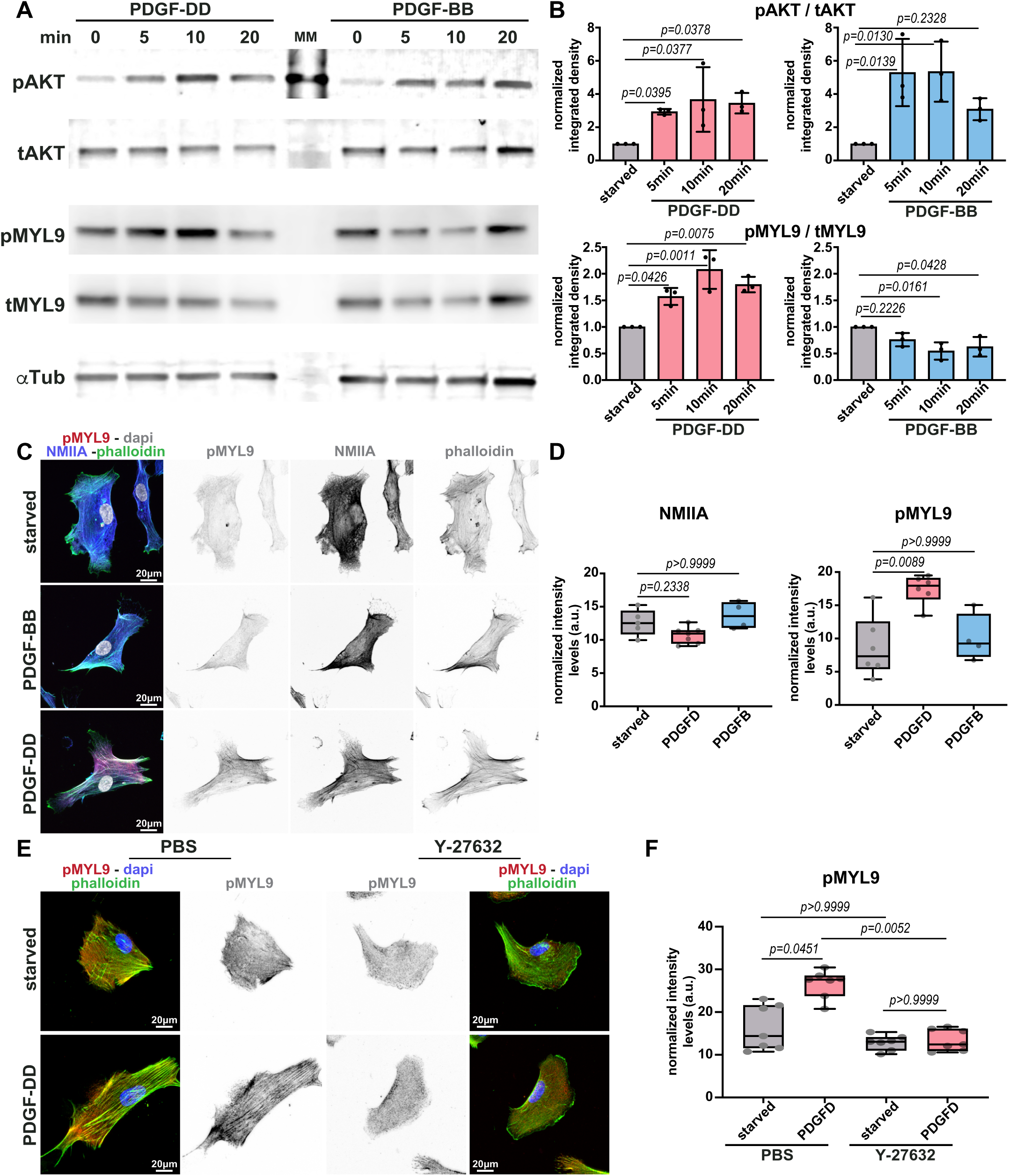
PDGFD, but not PDGFB, activates PC actomyosin contractility. **A)** Representative Western blot from human brain PCs (hBPCs) stimulated with PDGFD (40ng/mL) or PDGFB (40ng/mL) for 0, 5, 10, and 20 minutes. Protein levels of phosphorylated AKT (pAKT), total AKT (tAKT), phosphorylated MYL9 (pMYL9), total MYL9 (tMYL9) and α-tubulin (αTub) were assessed. **B)** Densitometric quantification of the pAKT/tAKT (top) and pMYL9/tMYL9 (bottom) ratios upon PDGF-DD (left) or PDGF-BB (right) stimulation, normalised to the starved condition. Data are mean ±s.d. from N=3 independent experiments. **C)** Representative images of hBPCs, starved or stimulated for 20 minutes with PDGF-BB (40ng/mL) or PDGF-DD (40ng/mL), and stained for pMYL9 (red), NMIIA (blue), filamentous actin (phalloidin, green) and nuclei (DAPI, grey). **D)** Quantification of NMIIA (left) and pMYL9 (right) mean fluorescence intensity in starved, PDGF-DD- or PDGF-BB-stimulated hBPCs. Data are presented as box plots with whiskers, min to max, with median. Each dot represents one experiment. N=6 starved; N=7 PDGF-DD; N=4 PDGF-BB. **E)** Representative images of hBPCs pre-treated with PBS or with the ROCK inhibitor Y-27632 (10µM) (30 minutes) and then unstimulated or stimulated for 20 minutes with PDGF-DD, stained for pMYL9 (red), filamentous actin (phalloidin, green) and nuclei (DAPI, blue). **F)** Quantification of the pMYL9 signal in PBS- or Y-27632-treated hBPCs, starved or stimulated with PDGF-DD. Data are presented as box plots with whiskers, min to max, with median. Each dot represents one experiment. N=7 for all conditions.

These results also suggest that inhibition of ROCK may reduce NVT formation. To test this hypothesis, we first assessed whether inhibition of ROCK could mimic NMIIA depletion by testing its impact on PC migration *in vitro*. We performed a scratch-wound assay on a confluent monolayer of hBPCs and measured wound closure. As expected, Y-27632 treatment significantly reduced cell migration and wound closure when compared to control treatment (PBS), consistent with ROCK inhibition mimicking NMIIA inactivation *in vitro* (Fig.S10A,B). Next, we administered Y-27632 or PBS, through intraperitoneal injection, to OIR mice two days after return to normoxia, at P13, and sacrificed the animals 3 days after injection, at P16 (Fig.S10C). Consistent with findings reported previously [50, 51], ROCK inhibition did not significantly alter the size of the avascular area (Fig.S10D-F). Yet, remarkably, Y-27632-treated animals displayed a strong reduction in both the total area and the average size of NVTs (Fig.S10D-H). Similar to NMIIA depletion in PCs, the decrease in NVT formation was not associated with a lack or decrease in NG2-positive PC coverage (Fig.S10I). Collectively, these findings strengthen the notion that pathological NVT formation depends on PDGFD– ROCK–NMIIA-driven actomyosin contractility in PCs, and suggest that targeting this pathway may be a promising therapeutic strategy for retinopathies.

### PDGFD drives NVT formation and PC actomyosin activation in vivo

Based on these results, we hypothesised that PDGFD loss-of-function would reduce NVT formation in mice. *Pdgfd* KO mice are viable and fertile and display a mild vascular phenotype [52]. Yet, the function of PDGFD in retinopathies has not been assessed. To test our hypothesis, littermate *Pdgfd*^wt/wt^ and *Pdgfd*^-/-^ mouse pups were subjected to the OIR protocol, and animals were sacrificed at P17 (Fig.7A). Because *Pdgfd*^-/-^ animals are constitutive knockouts, we first verified that the extent of hyperoxia-induced vaso-obliteration at P13 was indistinguishable between *Pdgfd*^wt/wt^ and *Pdgfd*^-/-^ retinas (Fig.S11A,B). At P17, *Pdgfd*^-/-^ animals showed a decreased avascular area and a significant reduction in both the total NVT area and the average size of individual NVTs (Fig.7B-F). Similar to NMIIA depletion or ROCK inhibition, the decrease in NVT formation was not associated with a loss of PC coverage (Fig.7G). Importantly, and in agreement with our *in vitro* data, NVTs in *Pdgfd*^-/-^ retinas displayed significantly reduced levels of both NMIIA and pMyl9 compared with NVTs of littermate controls (Fig.7H-J), directly linking PDGFD to the activation of the actomyosin machinery *in vivo*.

**Figure 7.**
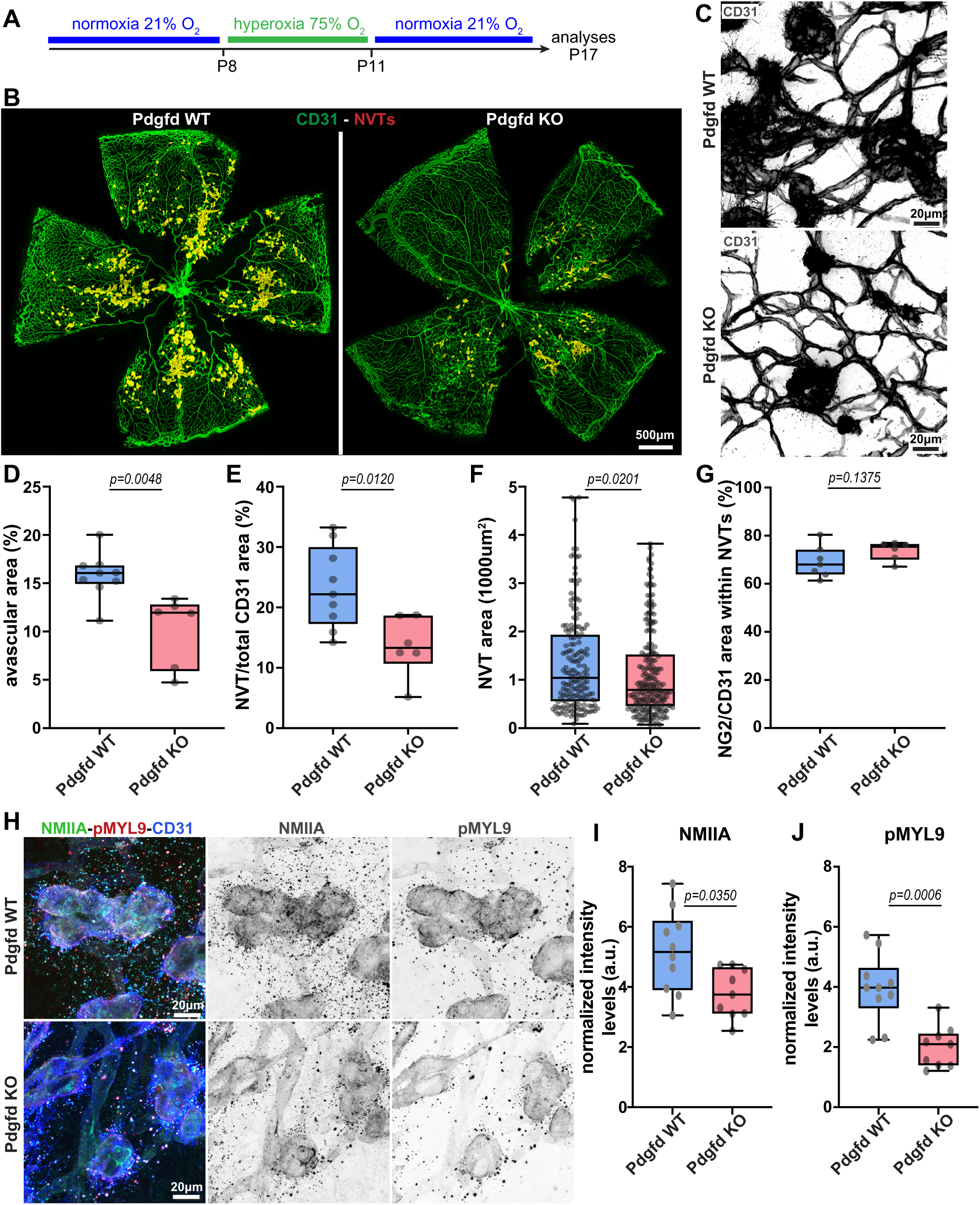
PDGFD drives NVT formation and PC actomyosin activation in vivo. **A)** Experimental timeline for the OIR protocol in Pdgfd^wt/wt^ (Pdgfd WT) and Pdgfd^-/-^ (Pdgfd KO) animals. **B)** Representative images of P17 OIR retinas from Pdgfd WT and Pdgfd KO mice stained for CD31 (green). Neovascular tufts (NVTs) were manually delineated (red) and superimposed on CD31 staining. **C)** Higher-magnification images of NVTs stained for CD31 (grey) from Pdgfd WT and Pdgfd KO mice. **D)** Quantification of the percentage of avascular area in P17 OIR retinas from Pdgfd WT and Pdgfd KO mice. Each dot represents one mouse retina. N=9 Pdgfd WT, N=6 Pdgfd KO. **E)** Quantification of the total NVT burden per retina, calculated as the sum of all NVT areas divided by the whole-retina CD31+ area (NVT/CD31 areas), in P17 OIR retinas from Pdgfd WT and Pdgfd KO mice. Each dot represents one mouse retina. N=9 Pdgfd WT, N=6 Pdgfd KO. **F)** Quantification of individual NVT area in P17 OIR retinas from Pdgfd WT and Pdgfd KO mice. Each dot represents one NVT, from N=9 Pdgfd WT and N=6 Pdgfd KO retinas. **G)** Quantification of the percentage of PC coverage (NG2+) in individual NVTs (CD31+) of P17 OIR retinas from Pdgfd WT and Pdgfd KO mice. Each dot represents one retina, from N=7 Pdgfd WT and N=6 Pdgfd KO. **H)** Representative images of NVTs of P17 OIR retinas from Pdgfd WT and Pdgfd KO mice, stained for NMIIA (green), pMyl9 (red) and CD31 (blue); single channels are shown as inverted greyscale. **I, J)** Quantification of NMIIA (I) and pMyl9 (J) levels in NVTs of P17 OIR retinas from Pdgfd WT and Pdgfd KO mice. Each dot represents one NVT. N=3 Pdgfd WT, N=3 Pdgfd KO retinas. All data are presented as box plots with whiskers, min to max, with median from 2 independent litters.

We next asked which cells express *Pdgfd* in the retina. Interrogation of the OIR scRNA-seq datasets showed that *Pdgfd* is expressed at low levels and in a sparse manner across several retinal cell populations, and is more strongly detected in ECs than in PCs in both normoxia and OIR (Fig.S11C-F). To complement this analysis, we took advantage of the LacZ cassette knocked into the *Pdgfd* locus of the *Pdgfd*^-/-^ allele [52] and performed X-gal staining on retinas of *Pdgfd*^wt/wt^ and *Pdgfd*^-/-^ littermates. X-gal signal was specific to *Pdgfd*^-/-^ retinas and was detected both in cells closely associated with the vasculature and in cells distributed throughout the retinal parenchyma (Fig.S11G). Collectively, these findings support the conclusion that pathological NVT formation is mostly driven by PC-dependent actomyosin contractility, which is regulated, in part, by PDGFD activity.

## DISCUSSION

Retinopathies such as PDR and ROP are major causes of vision loss, characterised by vascular leakage, inflammation, and the formation of aberrant NVTs [2, 3, 28]. These pathological structures form in response to ischemia-induced angiogenic signalling and are supported by dysfunctional EC and MC behaviour. PCs play a crucial role in both normal vascular development [38] and pathological contexts [12, 13, 16]. Here, we uncovered a novel molecular and cellular mechanism that drives NVT formation. We showed that NVT-associated ECs and PCs have very high levels of NMIIA and phosphorylated MLC, which are linked with specific cell morphologies, including blebs and rounded shapes. These features have been classically associated with high cortical actomyosin contractility [53, 54]. In addition, we uncovered that PDGFD-induced actomyosin contractility in PCs promotes NVT development and enlargement with minimal impact on neovascularisation. We also highlighted biomechanics as a driving factor for these vascular malformations, a component often underappreciated and understudied in this field. Remarkably, we showed that decreasing actomyosin contractility in ECs or PCs had opposite effects on NVT formation and development. Whilst NMIIA loss in ECs promoted larger and more numerous NVTs, NMIIA inactivation in PCs led to a decrease in the number and size of NVTs. Given the proximity and connectivity between both cell types, this result suggests an interplay of force balance that drives NVTs, which likely feeds back with EC proliferation. Although we do not explore this link further in this study, it would be interesting to analyse the roles of basement membrane composition and stiffness, as well as blood pressure, as relevant factors regulating force distribution within the PC-EC unit, and to assess their relative contributions to NVT formation and EC and PC proliferation.

Several *in vivo* studies have reported that ROCK inhibition attenuates pathological angiogenesis in OIR. For example, fasudil or ripasudil, two ROCK inhibitors, significantly decrease the avascular area and NVT area in OIR mice [47, 48, 55]. Nonetheless, it remains unclear how ROCK treatment promoted better recovery after the OIR protocol. For instance, most studies suggested that its main effects were on regulating EC contractility, migration and vasodilation properties. Yet, our results clearly indicate the opposite effect on ECs, as impairment of actomyosin contractility in ECs generated increased avascular area and NVT area. Instead, the effects of ROCK inhibitors were better mimicked by NMIIA deletion in PCs. Given the systemic effects of Y-27632, these observations strongly suggest that ROCK-mediated contractility may be more important in PCs than in ECs or other cell types for the development of NVTs. Activation of myosin motor function has been linked to ROCK but also to MRCK activity [49]. Thus, it is conceivable that PC actomyosin contractility may be more reliant on ROCK than that of ECs. In addition, a single intraperitoneal injection of Y-27632 at P13 produces a transient and partial inhibition of ROCK, whereas genetic deletion of *Myh9* removes the protein for the whole duration of the neovascular phase, which also may account for the differences in outcomes. Taking this into account, our results also suggest that inhibition of actomyosin contractility should be considered over a particular time window of intervention, as prolonged inhibition may also have detrimental effects on sprouting angiogenesis and revascularisation of the mouse retina.

Importantly, we identified PDGFD as a main regulator of PC actomyosin contractility and NVT formation. PDGFD appears to act as a context-dependent modulator of pathological angiogenesis rather than as a general factor required for PC migration in physiological conditions. We showed that *Pdgfd* loss markedly suppressed avascular area and NVT area, while *Pdgfd* loss has minimal effects on vascular development [52]. The role of *Pdgfd* as mostly relevant in pathological conditions is consistent with recent research highlighting PDGFD as a risk factor for atherosclerosis and stroke [56, 57]. In this study, we have not identified the precise cellular origin of PDGFD secretion. Using the LacZ reporter present in the *Pdgfd*-null allele and the OIR scRNA-seq datasets, we found that *Pdgfd* is expressed at low abundance in multiple cell types, including ECs (Fig.S11C-G). Several reports also point towards an endothelial origin in a disease context [51, 56, 57]. This could mean that, in addition to PDGFB, which regulates PC migration and proliferation, ECs communicate with PCs via PDGFD to regulate their contractility under pathological conditions. This is in agreement with a previous report that showed that PDGFD is mostly expressed in lung immune cells in physiological conditions, but widely expressed, including ECs, in idiopathic pulmonary hypertension disease [58]. Remarkably, we demonstrated PDGFRβ differential activation when stimulated with either PDGFB or PDGFD. PDGFD activated the phosphorylation of MLC and activation of actomyosin contractility more robustly than PDGFB, whilst pro-mitogenic AKT activation was more responsive to PDGFB. How the same receptor could elicit diverse responses to two distinct agonists remains unclear. A previous study has identified that PDGFD can bind to neuropilin-1 (NRP1) and signal through PDGFRβ in PCs [59], which may constitute a basis for differential downstream activation. Further studies may unravel the specific molecular signalling events that mediate divergent PDGFRβ downstream events upon PDGFB or PDGFD stimulation. Interestingly, this concept also provides a strong rationale for therapeutic selectivity. If PDGFD is mainly produced in activated ECs in pathological conditions and triggers selective signalling downstream of PDGFRβ, targeting PDGFD may blunt pathological behaviours whilst sparing physiological angiogenesis. This link is further supported by recent data identifying PDGFD as a highly enriched protein in the vitreous of PDR patients [47]. However, our study also reveals that PDGFD is likely not the only regulator of PC contractility and motility. *Pdgfd* deletion phenocopied ROCK inhibition and PC NMIIA loss in NVT formation and development, yet its phenotype was milder than direct *Myh9* deletion in PCs. Moreover, *Myh9* deletion strongly affects PC migration during development, whilst *Pdgfd* deletion has very minor defects in retinal angiogenesis. Therefore, further research aiming at characterising the additional factors contributing to PC contractility is required to fully harness the therapeutic potential of modulating this clinically relevant cellular activity.

Overall, our study supports a model in which NMIIA orchestrates the coordinated behaviour of ECs and PCs to maintain vascular structure and function during both physiological and pathological angiogenesis. We suggest that PDGFB-PDGFRβ signalling activates NCK and SRF transcriptional activity [13] in PCs. This signalling pathway likely facilitates the expression of actin-related genes, including Myh9, governing the migration of PCs, contributing to effective vessel stabilisation during developmental angiogenesis. In pathological conditions, PDGFD increases actomyosin activity through ROCK, leading to excessive PC contractility and NVT development.

In summary, our work establishes PDGFD and PC-derived contractility as key regulators of NVT formation and development in the mouse OIR model. Identifying specific mechanisms regulating PC actomyosin contractility, whilst leaving other cells, in particular ECs, untouched, holds promise for identifying therapeutic targets in proliferative retinopathies and other angiogenesis-driven disorders.

## MATERIALS AND METHODS

### Ethical statement

All animal experiments carried out in this work were performed in compliance with the relevant laws and guidelines that apply to the Gulbenkian Institute for Molecular Medicine (GiMM), Faculty of Medicine, University of Lisbon, Portugal. Animal procedures were approved by the local animal ethics committee and conducted with the permission of the Direção-Geral da Alimentação e Veterinária (DGAV), Portugal (0421/000/000/2016 and 0421/000/000/2021). Housing and procedures involving *Pdgfd^-/-^* mice were in accordance with Swedish legislation and approved by the local animal ethics committee (permit no. 11766-24).

### Bioinformatic analysis

Public mouse retinal single-cell RNA-sequencing (scRNA-seq) datasets from oxygen-induced retinopathy (OIR) and normoxic conditions (GEO accession numbers GSE150703 and GSE216676) were analysed using Seurat (v5.4.0). Cells with fewer than 200 detected genes, high mitochondrial gene expression, or excessive UMI counts were excluded during quality control. Potential doublets were identified using DoubletFinder (v2.0.4) and excluded from subsequent analyses. Each dataset was initially normalised and variance-stabilised using the SCTransform function. Principal component analysis (PCA), shared nearest-neighbour (SNN) graph construction, and clustering were subsequently performed, and Uniform Manifold Approximation and Projection (UMAP) was used for visualisation. ECs and PCs were selected from each dataset based on the expression of canonical marker genes (Cldn5, Pecam1, Cdh5, Pdgfrb, Rgs5, and Kcnj8). The selected populations were then merged, and the RNA assay was split into dataset-specific layers. Normalisation was performed using Seurat’s LogNormalize method, followed by identification of highly variable features using FindVariableFeatures function. The data were then scaled using ScaleData function, followed by PCA. Integration was performed using Seurat’s IntegrateLayers function with canonical correlation analysis (CCA) to account for possible dataset-associated batch effects. The integrated data were used for SNN graph construction and clustering, with UMAP for visualisation. Clusters were annotated as ECs, PCs, or proliferative subpopulations based on the expression of cell-type and proliferation markers, including Cldn5, Pecam1, Rgs5, Kcnj8, Mki67, and Top2a. Average gene expression was used to assess Myh and Myl gene expression across cell types and experimental conditions. Heatmaps were generated using ComplexHeatmap (v2.22.0).

### Mice and inducible genetic experiments

In this study, we used the following mouse strains: Myh9 floxed [60]; Rosa26^mTmG^ [34]; Rosa26^GNRep^ [43]; Cdh5(PAC)-Cre^ERT2^ [61]; Pdgfrβ-Cre^ERT2^ [62]; and Pdgfd^-/-^ [52] strains. All mice strains, except Pdgfd^-/-^, were maintained at GiMM under standard husbandry conditions (under specific pathogen-free conditions and kept in individually ventilated cages) and under national regulations. Pdgfd^-/-^ mice were maintained at the Karolinska Institute. The strains have a mixed background, and the intercrosses between the different mouse strains generated double and triple transgenic mouse strains. Postnatal gene deletion in newborn mice was induced via intraperitoneal (i.p.) injection of 50 µg 4OH-Tamoxifen (Sigma, #T5648), dissolved in ethanol-peanut oil (#P2144) at 1mg/ml on postnatal days (P) 1 to P3. In OIR experiments, animals were injected via i.p. with 200 µg of 4OH-Tamoxifen during 3 consecutive days between P11 and P13 in the Pdgfrβ-Cre^ERT2^ strain and with 100 µg at P11 and P13 for the Cdh5(PAC)-Cre^ERT2^ line. In our study, we shortened the hyperoxia exposure period to improve pup survival while still achieving substantial vascular regression and neovascular tuft formation. This protocol has been used in previous studies [7, 63, 64]. For the mosaic approach, pups were injected at P2 with 1ug or at P7 with 30 µL/g of a low dose of 4OH-Tamoxifen (0.04mg/ml). This approach was used to obtain individual mGFP+ ECs or MCs via stochastic recombination. All functional gene-deletion experiments used the full tamoxifen doses described above. In addition, animals were also injected via IP with a ROCK inhibitor, Y-27632 (MedChem, #HY-10071), at P13 (10mg/kg) and sacrificed at P16. Animals were sacrificed at several stages of development, and for each endpoint, the eyeballs were collected. For this study, both males and females were used without distinction for genetic experiments. As controls, Cre-negative littermates were used in the majority of the experiments. Cre-positive Myh9^wt/wt^ controls were used when performing tracing of R26^mTmG^ or R26^GNrep^ experiments.

### Oxygen-induced Retinopathy (OIR)

The OIR model was performed as previously described [28]. Neonatal pups and their mothers were kept in normoxic conditions (room air) from birth to P8 to allow normal vascular development. For 3 days from P8 to P11, the mice were placed under hyperoxic conditions of 75% O_2_ using the Biospherix A-Chamber (Biospherix). To minimize the stress of hyperoxia on the mothers, we used a modified protocol where the mothers were exposed to 22 hours of high oxygen and 2 hours of room air per day instead of continuous high oxygen exposure. This shortened hyperoxia protocol, first described by Kubota and colleagues [65], differs from the classical P7-P12 (5-day) protocol because it substantially reduces the mortality of nursing mothers and of the pups, while still producing extensive central vaso-obliteration and abundant NVT formation upon return to normoxia, which facilitates the quantification of tuft number and tuft size. After the hyperoxia period, the animals were kept for another 5 days, from P11 to P16, under normoxic conditions to induce the pathologic neovascular response. Later, animals were sacrificed, as well as their mothers, since these are no longer fertile after exposure to OIR, and thus should not be reused for breeding purposes. Animals were sacrificed by exposure to CO_2_ followed by cervical dislocation. Eyeballs were removed and fixed in 2% PFA. For Pdgfd mice, the protocol was 75% O_2_ from P8 to P11 and then normoxia until P17. The specific timing differences are related to the existing animal licenses for each institution, which differ slightly in their timings. All comparisons are made between littermate pups exposed to hyperoxia, processed and imaged in parallel.

### *In vivo* proliferation assay

To assess cell proliferation of ECs and MCs *in vivo*, thymidine analogue 5-Ethynyl-2′-deoxyuridine (EdU, Invitrogen, #10044) was prepared in PBS at 1mg/ml (for physiological conditions) and injected via IP (20μL/g of body weight) into P6 pups 4h before sacrifice. Retinas were prepared and stained using the Click-iT EdU Alexa Fluor-555 Imaging Kit (Invitrogen, #C10338) according to the manufacturer’s instructions and followed by an immunofluorescence protocol, as described below.

### Tissue preparation and immunofluorescence for whole-mount mouse retinas

Mice were sacrificed, and then the eyes were quickly removed using scissors and forceps, as described previously [66]. Then, they were fixed for 4 hours (or overnight) in 2% PFA in PBS at 4°C. After fixation, the eyeballs were washed in PBS and stored at 4°C until dissection, and retinas were dissected under a dissecting microscope. Next, retinas were incubated for 2 hours at RT in Claudio’s Blocking Buffer (CBB) (0.5% Triton X-100, 0.01% sodium deoxycholate, 0.02% sodium azide, 3% BSA and 1% fetal bovine serum (FBS) in PBS 1x pH=7.4) [67]. Next, they were incubated with primary antibodies (Table 1) in 1:1 PBS and CBB mixture at 4 °C overnight with gentle rocking. The next day, the primary antibody solution was removed, and the retinas were washed 3 times (30 min each) with PBST (PBS with 0.1% Triton X-100, (Sigma-Aldrich, #T8787)) and further incubated with secondary antibodies (Table 2) in 1:1 PBS and CBB mixture overnight at 4 °C or for 4 hours at RT with gentle rocking, protected from light. The next day, the retinas were washed 3 times (30 min each) with PBST and flat-mounted on glass slides using Vectashield mounting medium (Vector Laboratories, #H-1000). All mounted retina samples were stored protected from light at 4° C.

**Table 1.** List of primary antibodies used in immunofluorescent experiments.

| Antibody | Host | Dilution | Vendor/Catalog number |
| --- | --- | --- | --- |
| Desmin | Rabbit | 1/100 | Abcam (#ab15200) |
| VE-cadherin | Rat | 1/20 | BD Biosciences (#555289) |
| VE-cadherin | Mouse | 1/25 | Merck Millipore (#MABT134) |
| Collagen IV | Rabbit | 1/400 | Bio-Rad (#2150-1470) |
| NMIIA | Rabbit | 1/100 | Biolegend (#909801) |
| Cleaved Caspase-3 | Rabbit | 1/100 | Cell Signaling (#1679664S) |
| PDGFRbeta | Rat | 1/100 | ThermoFisher (#14-1402-82) |
| PECAM1 | Goat | 1/200 | R&D (#AF3628) |
| NG2 | Rabbit | 1/100 | Merck Millipore (#AB5320) |
| αSMA | Mouse | 1/100 | Sigma (#C6198) |
| pHistone H3 (S10) | Rabbit | 1/75 | Cell Signaling (#3475) |
| ERG | Rabbit | 1/200 | Abcam (#ab92513) |
| TER-119 | Rat | 1/100 | R&D (#MAB11259) |
| ICAM2 | Rat | 1/100 | BD Biosciences(#553326) |
| mCherry | Rat | 1/100 | ThermoFisher (#M11240) |
| GFP-FITC | Goat | 1/400 | Abcam (#ab6662) |
| pMLC2 | Rabbit | 1/200 | Cell Signaling (#3674) |
| pMYL9 | Mouse | 1/500 | ThermoFisher (#MA5-15163) |
| NMIIA | Rabbit | 1/400 | Sigma (#M8064) |
| CD13 | Rat | 1/100 | Bio-Rad (#MCA2183) |

**Table 2.** List of secondary antibodies used in immunofluorescent experiments.

| Antibody | Host | Dilution | Vendor/Catalog number |
| --- | --- | --- | --- |
| Phalloidin Alexa 568 | N/A | 1/400 | ThermoFisher (#A12380) |
| Phalloidin Alexa 647 | N/A | 1/400 | ThermoFisher (#A22287) |
| Phalloidin Alexa 488 | N/A | 1/400 | ThermoFisher (#A12379) |
| Anti-goat Alexa 647 | Donkey | 1/400 | ThermoFisher (#A21447) |
| Anti-rabbit Alexa 568 | Donkey | 1/400 | ThermoFisher (#A10042) |
| Anti-rabbit Alexa 488 | Donkey | 1/400 | ThermoFisher (#A21206) |
| Anti-mouse Alexa 488 | Donkey | 1/400 | ThermoFisher (#A21202) |
| Anti-rat Alexa 488 | Donkey | 1/400 | ThermoFisher (#A21208) |
| Anti-rabbit Alexa 647 | Donkey | 1/400 | ThermoFisher (#A31573) |
| Anti-goat Alexa 555 | Donkey | 1/400 | ThermoFisher (#A21432) |
| Anti-goat Alexa 488 | Donkey | 1/400 | ThermoFisher (#A11055) |
| Anti-mouse Alexa 555 | Donkey | 1/400 | ThermoFisher (#A31570) |

### Image acquisitions and morphometric analysis of retinal vasculature

Retina images were obtained from confocal microscope (Zeiss 880 and 980) equipped with the Zen black software with either an EC Plan-Neofluar 10x NA 0.30 dry, a Plan Apochromat 20x NA 0.80 dry, a C-Apochromat 40x NA 1.10 water or a Plan-Apochromat 63x NA 1.40 oil objectives. Tile scans covering the whole vascular area of the stained retinas were acquired on a spinning disk microscope (Zeiss Cell Observer SD) equipped with EC Plan-Neofluar 10x NA 0.30 dry, a Plan Apochromat 20x NA 0.80 dry, a Plan-Apochromat 40x NA 1.10 water or a Plan-Apochromat 63x NA 1.40 oil objectives and were stitched with the Zeiss blue software. In general, several fields of view from each animal (at least 4) were imaged and averaged. The number of animals in each group is indicated in the respective figures. Quantifications were performed with ImageJ (http://fiji.sc/) and IMARIS 9.8 (Oxford Instruments, https://imaris.oxinst.com/) software. In the mouse retina, the two regions were defined as: i) SF – region with 300µm of length from the periphery of the retina to the optic nerve and ii) RP – region closer to the optic nerve. The following vascular parameters were analyzed: radial expansion (the migratory length of the vascular plexus) was measured as distance from the optic nerve to the periphery of the vascular front; the number of tip cells was manually counted, and the values were normalized by the SF length; endothelial filopodia number at the sprouting front were counted manually and normalized to the tip cell number; vessel density was calculated as the ratio between vessel area and total area of the petal; vascularized area was measured as CD31+ area divided by total area; EC density, the number of ERG+ nuclei was counted manually and normalized by the vascularized area for each petal (two petals for each retina); MC migration *in vivo*, was measured as the distance between the ON and each GFP+ MC; cell proliferation – For EC proliferation, the number of proliferative ECs (double positive cells for ERG+ nuclei and EdU+) was manually counted and normalized to ERG+ cells. For MC proliferation, the number of proliferative MCs (EdU+, NG2+) was normalised to the NG2 area of the entire petal. The assessment of proliferation in the SF and in the RP was also performed and quantified as shown previously [68]. Collagen empty sleeves were counted manually as collagen IV+ area subtracted by the ICAM2+ vessel area, leaving only retracted vessels [35]; apoptotic ECs and MCs were manually counted as Casp3+ cells in PC (NG2+) or vessel (CD31+) areas; PC coverage was quantified as the specific marker (NG2, α-SMA, desmin and PDGFRβ)-positive area in the capillary areas and normalized to the vascularized area (CD31 area); extravasation of red blood cells (RBCs) was determined by manual selection of RBC area outside the vessel and then normalized to the total vessel area; MC Polarity Index was analysed in Rosa26^GNrep^ retinas, stained for GFP (nuclei), mCherry (golgi apparatus) and PCs (NG2). We adopted the analysis described previously [33, 44]. Briefly, we used PolNet as a graphical user interface to manually draw cell polarity vectors (nuclei to Golgi) for each mural cell in the network based on the GFP-mCherry image. Polarity index (PI) was then calculated using the angle that each Nucleus-to-Golgi vector makes with the SF edge, defined as a line between the two most outward vascular sprouts. The PI was calculated using consecutive Matlab scripts generated in the lab, and it varies between -1 (backward polarisation – away from the SF) and 1 (forward polarisation – towards the free edge of the SF), where 0 means random polarisation. The NVT and avascular areas of retinas were measured by using the “polygon selection” tool and normalised to the total vascularized area. The NVT burden is reported in two complementary ways: “NVT/total CD31 area (%)” corresponds to the sum of all NVT areas of a retina divided by the total CD31+ (whole-retina) vascular area of that same retina, whereas “NVT area” corresponds to the area of each tuft. NMIIA, pMLC2 and pMYL9 levels in NVTs versus non-NVT vessels were quantified on maximum-intensity projections by manually delineating individual NVTs and an equivalent number of non-tuft capillary fields within the same retina, measuring the mean fluorescence intensity inside the vessel mask and normalising to the area of each object. NMIIA levels in single PCs or ECs were quantified as a percentage of recombination for each stage in the mouse retina. Using Imaris, we masked individual GFP+ cells to quantify the GFP+ area of each cell and then measured NMIIA fluorescence intensity in the Myh9 channel. Because NMIIA is ubiquitously expressed in the retina, only the NMIIA signal contained within the three-dimensional surface of each isolated recombined cell was retained. Given the resolution limits of the technique and the close relationship between ECs and PCs, the masks for ECs or PCs do contain signal from the neighbouring cells. 3D visualisation videos of segmented mGFP+ cells was performed using IMARIS 9.8 software.

### RNA extraction, cDNA synthesis and RT-qPCR

For RNA extraction from mouse retinas, the pair of retinas from an animal was dissected in ice-cold fresh PBS and immediately snap-frozen in liquid nitrogen. Retinas were lysed in RLT buffer from the RNeasy Micro Kit (Qiagen, #74004), vortexed vigorously and further homogenised using 30G syringe needles. Lysates were centrifuged to clear precipitates, and the supernatant was used for RNA extraction as described by the manufacturer’s protocol, including in-column DNase I treatment. RNA concentration was quantified using NanoDrop 2000 (ThermoFisher) and adjusted equally for complementary DNA (cDNA) synthesis (Superscript IV First-Strand Synthesis System, Invitrogen, #18091050). The cDNA samples were then diluted in RNAse/DNAse-free water for the subsequent real-time quantitative PCR (RT-qPCR) reactions. RT-qPCR was performed using a 7500 Fast Real-Time PCR System (Applied Biosystems) with Power SYBR Green PCR Master Mix (ThermoFisher, #4368702) following the standard program of the previously mentioned system. For each reaction, 5μL of cDNA was combined with 10μL of Power SYBR Green PCR Master Mix, 4.5μL of RNAse/DNAse-free water and 0.5μL of 4μM primer pool (forward+reverse) (Table 3) in a MicroAmp Fast Optical 96-well Reaction Plate (Applied Biosystems). The expression levels of each sample duplicate were then normalised to GAPDH, and the Livak method was used to calculate relative alterations in gene expression.

**Table 3.**
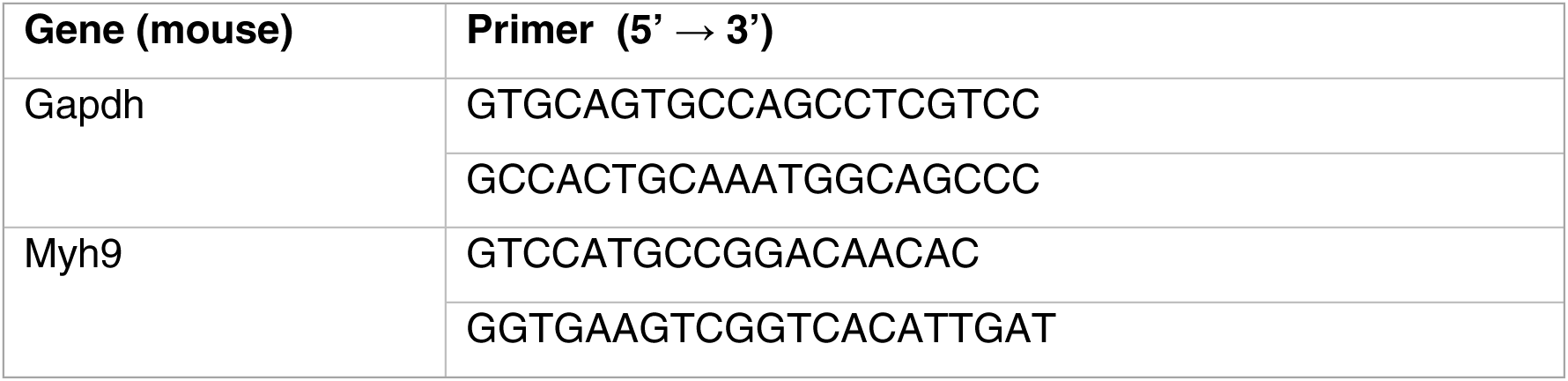
List of the mouse primers used in the qPCR.

### Isolation and culture of primary mouse brain mural cells

Primary mouse brain mural cells (mBMCs) were isolated from 6- to 8-week-old Myh9 MC-iKO::R26^mTmG^ animals. The isolation procedure was performed as previously described by Tiggers et al. [69]. Briefly, brains from 3 or more mice were dissected and placed in ice-cold PBS 1x (137mM NaCl, 2.7mM KCl, 4.3mM Na^2^HPO^4^, 1.47mM KH^2^PO^4^, pH 7.4) containing 1% Pen/Strep. Using a blade, the olfactory bulb, cerebellum and medulla were removed, and the remaining cortical tissues were pooled and rigorously minced. Then, the brains were washed using PBS and centrifuged for 5 min at 300g at room temperature (RT). The tissue pellet was homogenised in an enzymatic mixture by slow trituration, passing it 10 times through an 18G needle, followed by passing it 10 times through a 20G needle and finally through a 21G needle. The enzymatic digestion mixture contains DNase I (1mg/mL, Sigma, #10104159001) and collagenase II (1mg/ml, Worthington Biochemical, # LS004176) and was incubated at 37 °C for 70 min. After incubation, the digested tissue was then resuspended in 1.7x v/v of 22% bovine serum albumin (BSA) (NZYTech, # MB04602)/PBS solution and centrifuged at 1300g for 10 min to separate the cell content in the pellet from the myelin ring in the upper phase. The supernatant was removed completely by aspiration and the cell pellet was then washed in EGM2 medium (Lonza, #CC-3162 and #CC-3121) followed by centrifugation at 300g for 5 min. After aspiration of the supernatant, the pellet containing single cells was resuspended in 2 mL EGM2 medium and seeded in 1 well of a 6-well plate coated with 0.2% collagen type I (Corning, #CB354249). Plated cells were incubated at 37 °C and 5% CO^2^ for 20 hours in a humidified incubator to ensure attachment of viable cells to the plate. After incubation, the cells were carefully washed 3 times with PBS. Finally, fresh EGM-2 medium was added to each well, and the cells were cultured until reaching confluence. Then, the cells were trypsinised (TrypLE™ Express Enzyme (1X) - ThermoFisher, #12605028), pooled and further cultured in a T25 cell culture flask. The established cell culture, containing ECs, fibroblasts and PCs, was maintained in EGM-2 medium until the second passage. The mBMC culture was established by growing the cells in PC-medium (ScienCell, #1201) from the third passage onwards, which especially promotes MC growth. In order to increase the purity of our culture, after the third passage, the cells were incubated with Dynabeads (Invitrogen, # 11035) coated with the PDGFRβ antibody. Briefly, 80µL beads/T25 flask were added to a sterile Eppendorf, and they were washed 5 times with 0.5% BSA in PBS and mixed well by pipetting up and down. Then, the Eppendorf was placed in a magnet rack, and the beads were attached to the wall adjacent to the magnet. Next, a PDGFRβ antibody was added (8µL/T25 flask) and incubated for 1 hour at RT covered from light. During this period, mBMCs in culture were trypsinised and centrifuged, as explained previously, and then the pellet was resuspended in 100µl of PBS/0.5% BSA. After 1 hour of incubating beads with the antibodies, the bead-antibody solution was put into 100µL of cell suspension and incubated for 1 hour at RT and protected from light. After this incubation, the beads were washed 5 times with 0.5% BSA in PBS and the Eppendorf tube was placed into the magnet rack to separate the beads. Finally, the cell suspension was resuspended in PC-medium and added to a previously coated T25-flask. On the next day, the medium was changed, and the cells were kept until they reached confluence.

### Cre activation of mBMCs *in vitro*

To promote Cre-mediated recombination, 2x10^5^ cells/6-well or 5x10^4^ cells/24-well of mBMCs were seeded on 0.2% collagen type I-coated plates and cultured overnight. The next day, cells were washed with PBS, and then, 4OH-Tam was added to the wells (final concentration 5μM) or, as a control, ethanol was added to the PC medium. The cells were incubated for 72 hours, and after that time, they were directly used for experiments. Furthermore, the protein level was determined by Western blot analyses.

### Immunofluorescence in cells and image acquisition

To perform immunofluorescence, cells were fixed in 4% paraformaldehyde (PFA) (#15710, Electron Microscopy Sciences) supplemented with 1M MgCl_2_ and 1M CaCl_2_ (1μL/2mL) in PBS for 10 min at RT. Then, cells were washed with PBS 1x to remove the remaining PFA, followed by blocking and permeabilisation of cells with a blocking solution containing 3% BSA in PBS-T (PBS with 0.1% Triton X-100) for 30 min at RT. After, cells were incubated with primary antibodies for 2 hours at RT diluted in the blocking solution (Table 1) and washed 3 x 15min in PBS-T. Next, cells were incubated in blocking solution containing the secondary fluorophore-conjugated antibodies for 1h at RT in the dark (Table 2), followed by 3 washes of 15min in PBS-T. Finally, cells were incubated with 1/1000 (v/v) DAPI/PBS (Molecular Probes by Life Technologies, #D1306) for 5 min in the dark. Ibidi slides were mounted in 20% glycerol in PBS into the channel, and the coverslips were mounted in Mowiol-DABCO. The image acquisition was performed in the Zeiss Cell Observer SD with a Plan Apochromat 20x NA 0.80 dry and in the confocal Zeiss 880 with a Plan-Apochromat 63x NA 1.40 oil objective. Both microscopes were equipped with the Zen software.

### *In vitro* proliferation assay

To assess cell proliferation of mBMCs *in vitro*, thymidine analogue 5-ethynyl-2′-deoxyuridine (EdU, Invitrogen, #10044) was prepared at 10mM and added to a final concentration of 20μM to the cell medium 2 hours before fixation. Coverslips were prepared and stained using the Click-iT EdU Alexa Fluor-555 Imaging Kit (Invitrogen, #C10338) according to the manufacturer’s instructions. The number of cells in proliferation was quantified as EdU+Dapi+ cells and then normalized to the total number of cells in each coverslip.

### Protein extraction and western blotting

mBMCs (control (EtOH) and *Myh9* knockout (*Myh9*^KO^) (4OH-Tam) were grown until they reached 90% confluence on 6-well cell culture plates. Similarly, overnight starved hBPCs (Innoprot, #P10363) were stimulated with human PDGFD (40ng/mL, Genscript, #Z03195) and human PDGFB (40ng/mL, ThermoFisher, #100-14B) for 0, 5, 10 and 20 minutes. Cell lysis was performed with a cold Lysis buffer (150mM NaCl, 1mM ethylenediamine tetracetic acid (EDTA), 50mM Tris-HCl pH 7.4, 1% Triton X-100 (Sigma, #T8787), supplemented with 1mM DTT (Sigma, #D0632) and proteinase and phosphatase inhibitor (1:100) (ThermoFisher, # 78446) in H_2_O, followed by a 10 min centrifugation at 10000g at 4° C. The supernatant containing proteins was then transferred into a fresh tube, and the protein concentration was determined by the Pierce BCA Protein Assay Kit (ThermoFisher, #23227). Protein lysates were stored at -20 °C until used. 20-50µg of protein was incubated in lysis buffer at 37 °C for 30 min, then SDS-PAGE was performed, and subsequently the separated samples were transferred onto a 0.22μm polyvinylidene difluoride transfer membrane using a Bio-Rad transfer system for 2 hours at 250mA. Then, the membranes were washed in TBST buffer (138mM NaCl, 20mM Tris, 0.1% Tween-20 in H_2_O, pH 7.4). The membrane was subsequently blocked in a blocking buffer containing 3% BSA in TBST for 1 hour at RT. Next, membranes were incubated with primary antibodies (Table 4) diluted in the blocking buffer overnight at 4 °C. On the following day, the membranes were washed 3 times with TBST and subsequently incubated with corresponding horseradish peroxidase (HRP)-coupled secondary antibodies (Table 5) diluted in the blocking buffer for 1 hour at RT. Finally, membranes were washed three times with TBST. Imaging of membranes was performed by the addition of chemiluminescent HRP substrate, ECL Prime Western Blotting Detection Reagent (GE Healthcare, #RPN2232) or by ECL Select Western Blotting Detection Reagent (GE Healthcare, #RPN2235), and the membranes were revealed in Amersham 680 or 800 (GE Healthcare). When IRDye secondary antibodies were used, the membranes were protected from light, the antibody incubations were done in TBST, and one final wash in TBS for 5 min was added to remove the detergent before exposure using the Odyssey Infrared Scanner and the Image Studio software. Phosphorylated and total forms of the same protein were always detected on the same membrane. Densitometric quantification for phospho-proteins was performed in Fiji on non-saturated exposures: for each lane, the integrated density of the band of interest was measured after local background subtraction, normalised to the total protein of the same lane, and finally normalised to the unstimulated condition of the same experiment.

**Table 4.** List of primary antibodies used in Western blots.

| Antibody | Host | Dilution | Vendor/Catalogue number |
| --- | --- | --- | --- |
| NMIIA | Rabbit | 1/1000 | Biolegend (#909801) |
| GFP | Goat | 1/5000 | Abcam (#ab6662) |
| β-actin | Mouse | 1/1000 | Sigma (#A5441) |
| pMYL9 | Mouse | 1/500 | ThermoFisher (#MA5-15163) |
| MYL9 | Rabbit | 1/1000 | Abcam (#ab64161) |
| pAKT | Rabbit | 1/5000 | Cell Signaling (#4060) |
| AKT | Rabbit | 1/2000 | Cell Signaling (#9272) |
| α-Tubulin | Mouse | 1/2000 | Sigma (#T6199) |

**Table 5.** List of secondary antibodies used in Western blots.

| Antibody | Host | Dilution | Vendor/Catalogue number |
| --- | --- | --- | --- |
| Anti-Rabbit IgG H&L (HRP) | Donkey | 1/10000 | Abcam (#ab6802) |
| Anti-Mouse IgG H&L (HRP) | Donkey | 1/10000 | Abcam (#ab6820) |
| Anti-Goat IgG H&L (HRP) | Donkey | 1/5000 | Abcam (#ab205723) |
| Anti-Rabbit IgG (IRDye 680) | Donkey | 1/5000 | LICORbio (#926-68073) |
| Anti-Goat IgG (IRDye 800) | Donkey | 1/2000 | LICORbio (#926-32214) |
| Anti-Mouse IgG (IRDye 800) | Donkey | 1/5000 | LICORbio (#926-32212) |

### Scratch-wound migration assay

*Myh9*^KO^ and the respective control of mBMCs were seeded to 12-well plates coated with 0.2% collagen, and the cells were cultured in PC medium. The Cre was activated with 4OH-Tam or, as a control, we added ETOH. Similarly, hBPCs treated with PBS as a control, or with a ROCK inhibitor (Y-27632, MedChem, #HY-10071, 10µM) were seeded onto µ-Slide 8-well slides (IBIDI, #80826). In both cases, on the day of the experiment, the wound was made with a yellow tip. The wells were washed with PBS, and fresh PC medium was added. The cells were imaged in the wound region during 14 or 16 hours at 37 °C using the Zeiss Cell Observer SD microscope with a Plan Neofluar 10x NA 0.30 dry objective. The wound area was determined by manually outlining the scratch in each well at each time point. The migrated area was calculated as the initial wound area minus the remaining wound area at the final time point. The cell velocity was quantified by single-cell tracking of cells near the scratch using the Chemotaxis and migration tool.

### **X-** gal staining

The Pdgfd KO allele carries a LacZ reporter cassette knocked into the Pdgfd locus [51], which allows Pdgfd-expressing cells to be visualised by β-galactosidase activity. Eyes from Pdgfd^wt/wt^ and Pdgfd^-/-^ littermates were fixed in 4% PFA for 1 h at room temperature and retinas were dissected and washed in rinse buffer (0.1 M phosphate buffer (pH 7.3), 2 mM MgCl_2_, 0.01% sodium deoxycholate, 0.02% Nonidet P-40) twice, followed by incubation in staining solution (0.1 M phosphate buffer (pH 7.3), 2 mM MgCl_2_, 0.01% sodium deoxycholate, 0.02% Nonidet P-40, 5 mM potassium ferricyanide, 5 mM potassium ferrocyanide, 1 mg ml^−1^ X-gal) for 3 h at 37 °C and then washed in rinse buffer 2X20 min at room temperature. Samples were then post-fixed in 4% PFA for 10 min before immunostaining with antibodies.

### Statistical analysis

Data were analysed with GraphPad Prism 8 software. Although no statistical methods were used to predetermine sample size, we aimed to have experimental groups of at least two different litters for all *in vivo* experiments, based on our expertise (minimum number to provide robust and reliable data). Measurements were taken from distinct samples, and statistical details of experiments are reported in the figures and figure legends. Sample size is reported in the figure legends. The biological replicate is defined as the number of cells, number of retinas, NVT images or animals, as stated in the figure legends. No inclusion/exclusion or randomisation criteria were used, and all analysed samples are included. Normality tests were performed to assess the data normality. Comparisons between two experimental groups were analysed with a two-sided unpaired parametric t-test or a Mann-Whitney test, depending on the data normality. Multiple comparisons between more than two experimental groups were assessed with one-way ANOVA. We considered the result significant when p<0.05. For all box plots: central line, median; +, mean; whiskers, min to max.

## Acknowledgments

We would like to express our appreciation to all members of the Vascular Biology and Disease Laboratory (previous and current) and the Fondation Leducq ATTRACT members for their active participation in extensive and helpful scientific discussions. Additionally, we acknowledge the staff from CBR and the technical support of the Bioimaging Platform of the Gulbenkian Institute for Molecular Medicine (GIMM), funded by PPBI-POCI-01-0145-FEDER-022122. Lastly, we would like to thank the dedicated staff from the animal facility of GIMM for their exceptional support in mouse husbandry.

## Sources of Funding

This work was supported by Fundação para a Ciência e a Tecnologia (FCT) (PTDC/MED-ANM/7695/2020; 2022.09228.PTDC, and UIDP/04378/2020), Fondation Leducq (17CVD03) and ‘la Caixa’ Foundation (LCF/PR/HR22/52420027; LCF/PR/HR24/52440027). Work by the L.J. laboratory was supported by the Swedish Research Council (2021-01210), the Swedish Cancer Society (222440 Pj, 254817 Pj), the Swedish Brain Foundation (FO2022-0055, FO2025-0238-HK-186), the Swedish Heart and Lung Foundation (20220354, 20250759), Grünwald Goglund stiftelse and Karolinska Institutet. A.P. was supported by a personal PhD fellowship from FCT (SFRH/BD/130635/2017 and COVID/BD/152399/2022). D.R. was supported by a personal PhD fellowship from FCT (2020.05174.BD). A.F. was supported by a postdoctoral researcher contract from Fundação para a Ciência e a Tecnologia (2021.04013.CEECIND).

## Author contributions

Conceptualization: A.P., A.M.F., C.A.F.

Methodology: A.P., D.R., M.P.S., Y.J., E.B.F., U.E., L.J., C.A.F.

Experimentation and data acquisition: A.P., D.R., M.P.S., Y.J., M.M., C.E., L.H.M., M.O., Y.C., L.J., A.M.F., C.A.F.

Data analysis: A.P., D.R., M.P.S., Y.J., M.M., L.J., A.M.F., C.A.F.

Project administration: L.J., C.A.F.

Writing – original draft: A.P., A.M.F., C.A.F.

Writing – review & editing: A.P., D.R., L.H.M., Y.J., E.B.F., U.E., L.J., A.M.F., C.A.F.

## Data Availability Statement

All data supporting the findings of this study are available within the article and its Supplementary Information. Source data for all graphs are available from the corresponding author on reasonable request. No new sequencing data were generated in this study.

## Competing interests

The authors declare that they have no competing interests.

## Abbreviations

αSMA -: alpha-smooth muscle actin
DR -: diabetic retinopathy
EC -: endothelial cell
MC -: mural cell
Myh -: myosin
NCK -: non-catalytic region of tyrosine kinase adaptor protein 1
NG2 -: neural/glial antigen 2
NMII -: non-muscle myosin II
MYL –: myosin light chain
NVT -: neovascular tuft
OIR -: oxygen-induced retinopathy
PC -: pericyte
PDGF -: platelet-derived growth factor
PDGFRꞵ -: platelet-derived growth factor receptor beta
PDR -: proliferative diabetic retinopathy
ROP -: retinopathy of prematurity
RP -: remodeling plexus
SF -: sprouting front
SMC -: smooth muscle cell
SRF -: serum response factor
TAM -: tamoxifen
VEGF -: vascular endothelial growth factor
vSMC -: vascular smooth muscle cell

## SUPPLEMENTAL FIGURE LEGENDS

**Figure S1.**
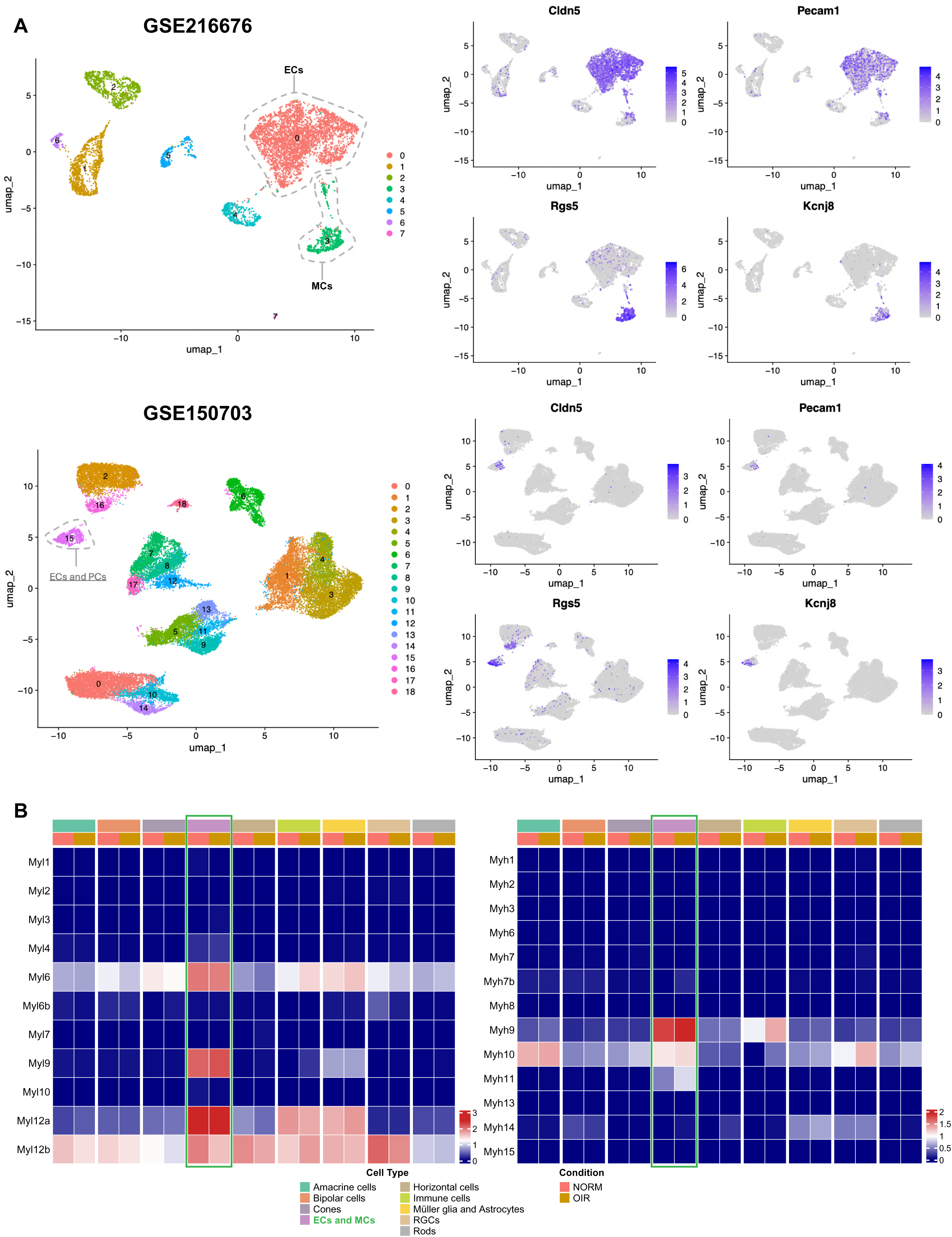
Myosin gene expression across retinal OIR scRNAseq datasets. **A)** UMAPs of EC and PC clusters. (left) UMAPs of GSE216676 (top) and GSE150703 (bottom) with EC and PC clusters highlighted by a discontinuous grey outline, and (right) feature plots showing the expression of canonical markers Cldn5, Pecam1 (ECs) and Rgs5, Kcnj8 (PCs) across clusters. **B)** Heatmaps of average expression levels of myosin light chain (Myl; left) and heavy chain (Myh; right) genes across identified clusters in GSE150703 under normoxia and OIR conditions.

**Figure S2.**
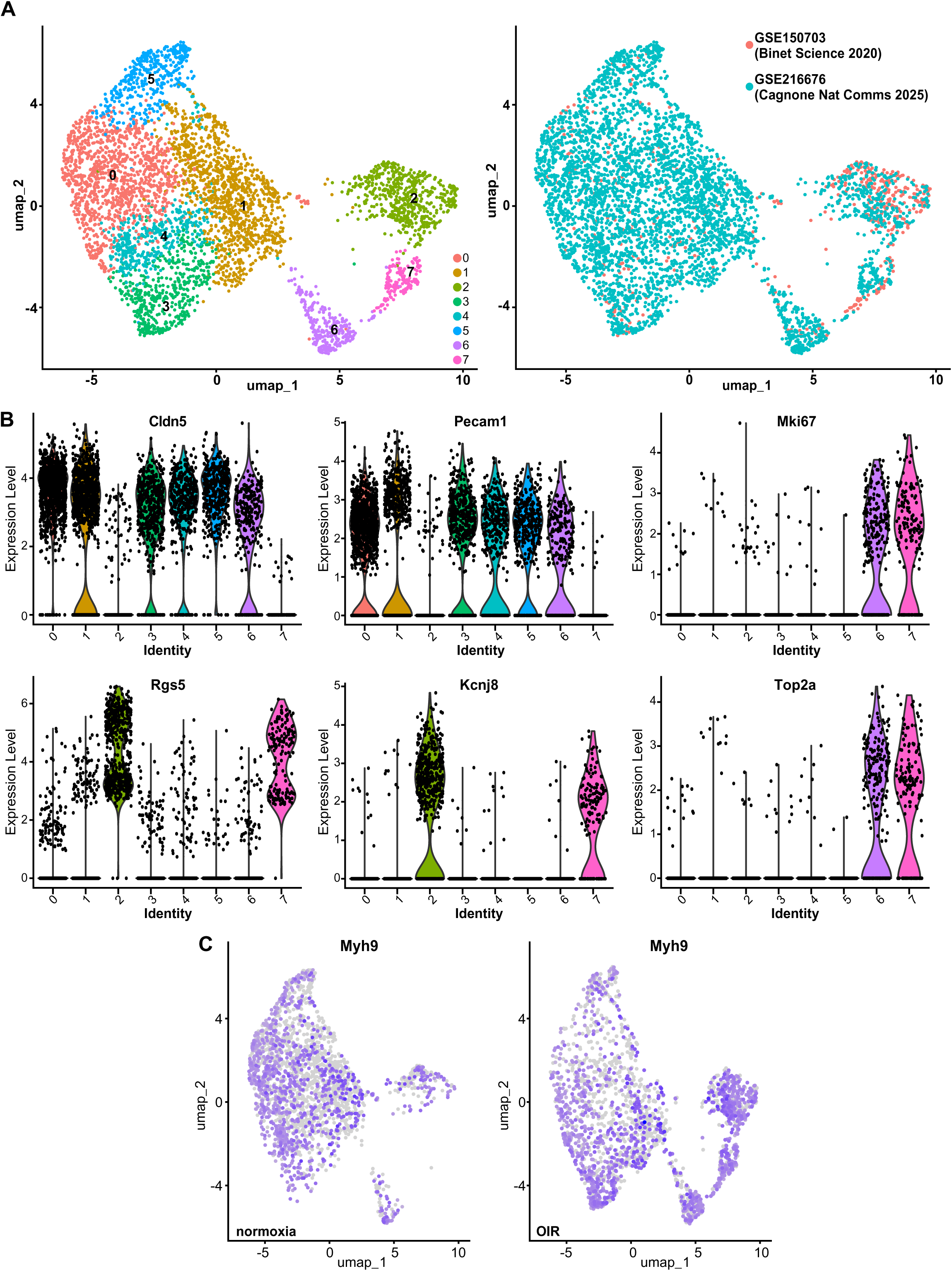
Integration and validation of vascular cell identities in scRNAseq datasets. **A)** UMAP integration of two independent retinal scRNAseq datasets. **B)** Violin plots showing expression of canonical EC and PC markers (Cldn5, Pecam1, Rgs5, Kcnj8, Top2a, Mki67) across identified clusters. **C)** UMAPs of EC and PC clusters showing expression of Myh9 in normoxia and OIR conditions.

**Figure S3.**
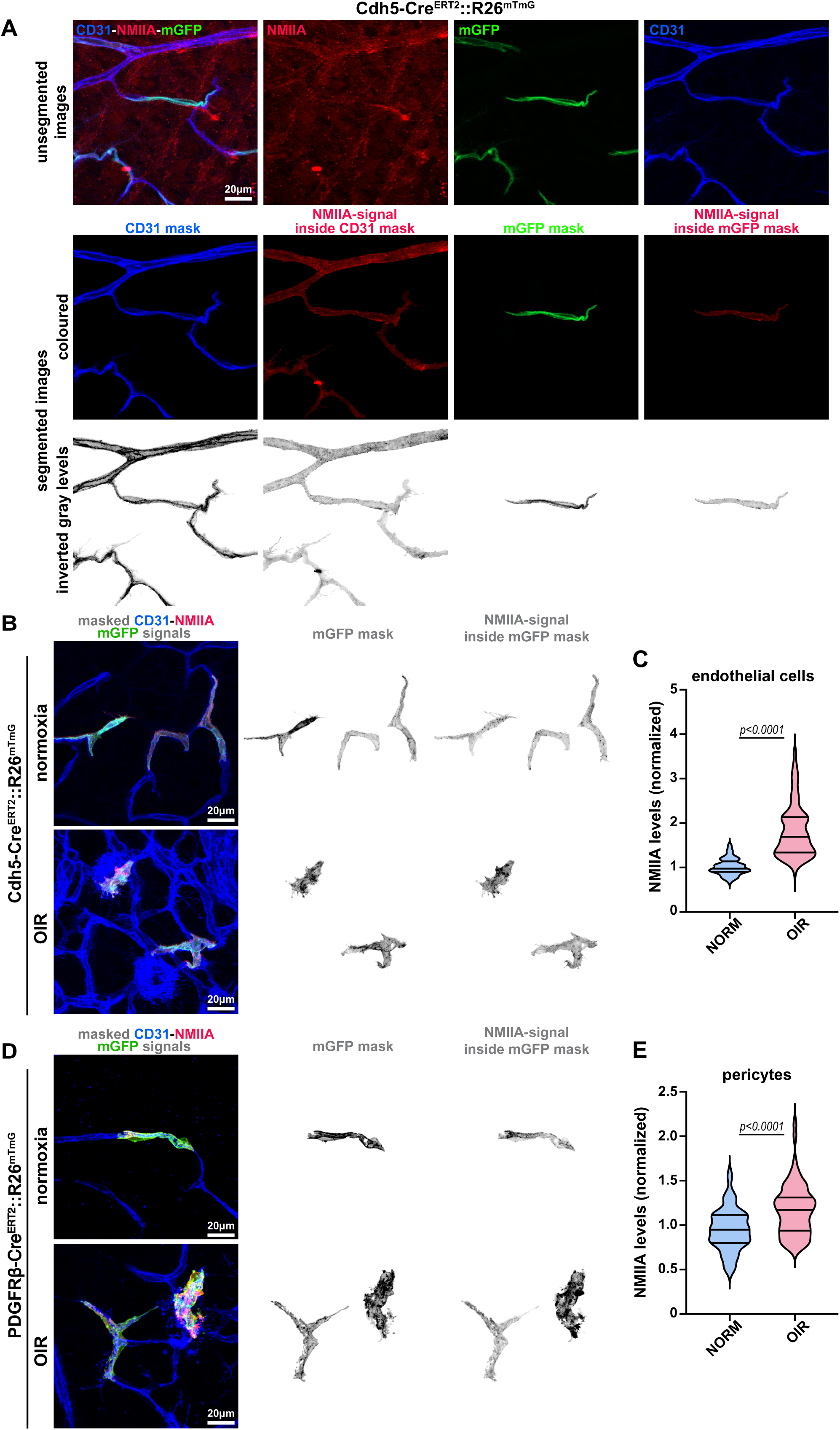
Single-cell quantification of NMIIA levels in ECs and PCs in normoxia and in OIR. **A)** Workflow used to quantify NMIIA in single recombined cells. Top row: unsegmented confocal images of a Cdh5^CreERT2^::R26^mTmG^ OIR retina showing the merge and the individual NMIIA (red), mGFP (green) and CD31 (blue) channels. Middle and bottom rows: the same field after segmentation, displayed in colour and as inverted greyscale, showing the CD31 mask, the NMIIA signal contained within the CD31 mask, the mGFP mask of a single recombined cell, and the NMIIA signal contained within that mGFP mask. See also Supplementary Video 1. **B)** Representative images of single mGFP+ ECs from Cdh5^CreERT2^::R26^mTmG^ retinas under normoxia or OIR, with masked images of blood vessels (CD31, blue, in overlay), masked single mGFP+ ECs (green on overlay, grey on inverted grey panels) and the NMIIA signal inside the mGFP mask (red on overlay, grey on inverted grey panels). **C)** Violin plot for the quantification of NMIIA levels, normalised to normoxia, in single ECs from retinas of animals exposed to normoxic (NORM) or OIR conditions. N=84 in NORM and N=194 in OIR. **D)** Representative images of single PCs of PDGFRβ^CreERT2^::R26^mTmG^ retinas under normoxia or OIR, with masked images of blood vessels (CD31, blue, in overlay), masked single mGFP+ PCs (green on overlay, grey on inverted grey panels) and the NMIIA signal inside the mGFP mask (red on overlay, grey on inverted grey panels). **E)** Violin plot for the quantification of NMIIA levels, normalised to normoxia, in single PCs. Each dot represents one MC. N=139 in NORM and N=133 in OIR. Data in C and E are presented as violin plots with median and quartiles, from at least 3 independent litters.

**Figure S4.**
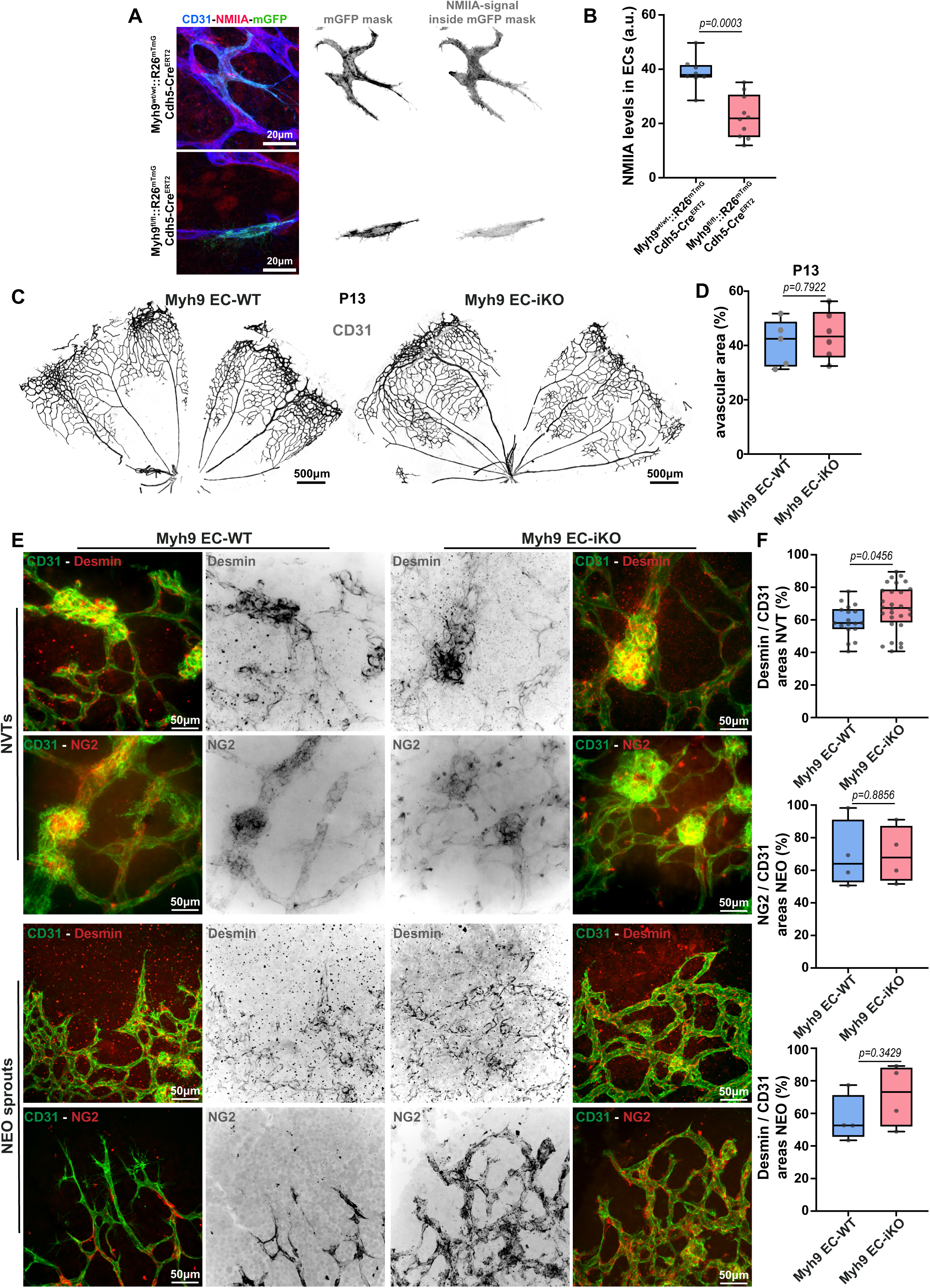
PC coverage in neovascular and neo-angiogenic regions after EC-specific deletion of Myh9. **A)** Representative images of NMIIA immunofluorescence in ECs of P16 OIR retinas from Myh9^wt/wt^::Cdh5-Cre^ERT2^::R26^mTmG^ and Myh9^fl/fl^::Cdh5-Cre^ERT2^::R26^mTmG^ mice. **B)** Quantification of NMIIA levels in ECs of P16 OIR retinas from Myh9^wt/wt^::Cdh5-Cre^ERT2^::R26^mTmG^ and Myh9^fl/fl^::Cdh5-Cre^ERT2^::R26^mTmG^. Each dot represents one retina. N=8 Myh9^wt/wt^::Cdh5-Cre^ERT2^::R26^mTmG^; N=10 Myh9^fl/fl^::Cdh5-Cre^ERT2^::R26^mTmG^. **C)** Representative images of P13 (post-hyperoxia) retinas from Myh9^EC-WT^ and Myh9^EC-iKO^ mice stained for CD31 (grey). **D)** Quantification of the percentage of avascular area at P13 in Myh9^EC-WT^ and Myh9^EC-iKO^ retinas. Each dot represents one mouse retina. N=5 mice for *Myh9*^EC-WT^ and N=6 mice for *Myh9*^EC-iKO^. **E)** Representative images of NVTs (upper panels) or neovascular sprouts (NEO) (bottom panels) of P16 OIR retinas from *Myh9*^EC-WT^ (left panels) and *Myh9*^EC-iKO^ (right panels) mice, stained for CD31 (green) and desmin (red and grey; top panel) or NG2 (red and grey; bottom panel). **F)** Quantification of the percentage of PC coverage (desmin) in individual NVTs, and of NG2 and desmin coverage in neovascular sprouts (CD31+) of P16 OIR retinas from *Myh9*^EC-WT^ and *Myh9*^EC-iKO^ mice. For NVTs, each dot represents one NVT. N=18 *Myh9*^EC-WT^; N=28 *Myh9*^EC-iKO^ NVTs. For NEO, each dot represents one retina. N=4 mice for *Myh9*^EC-WT^ and *Myh9*^EC-iKO^. All data are presented as box plots with whiskers, min to max, with median from at least 3 independent litters.

**Figure S5.**
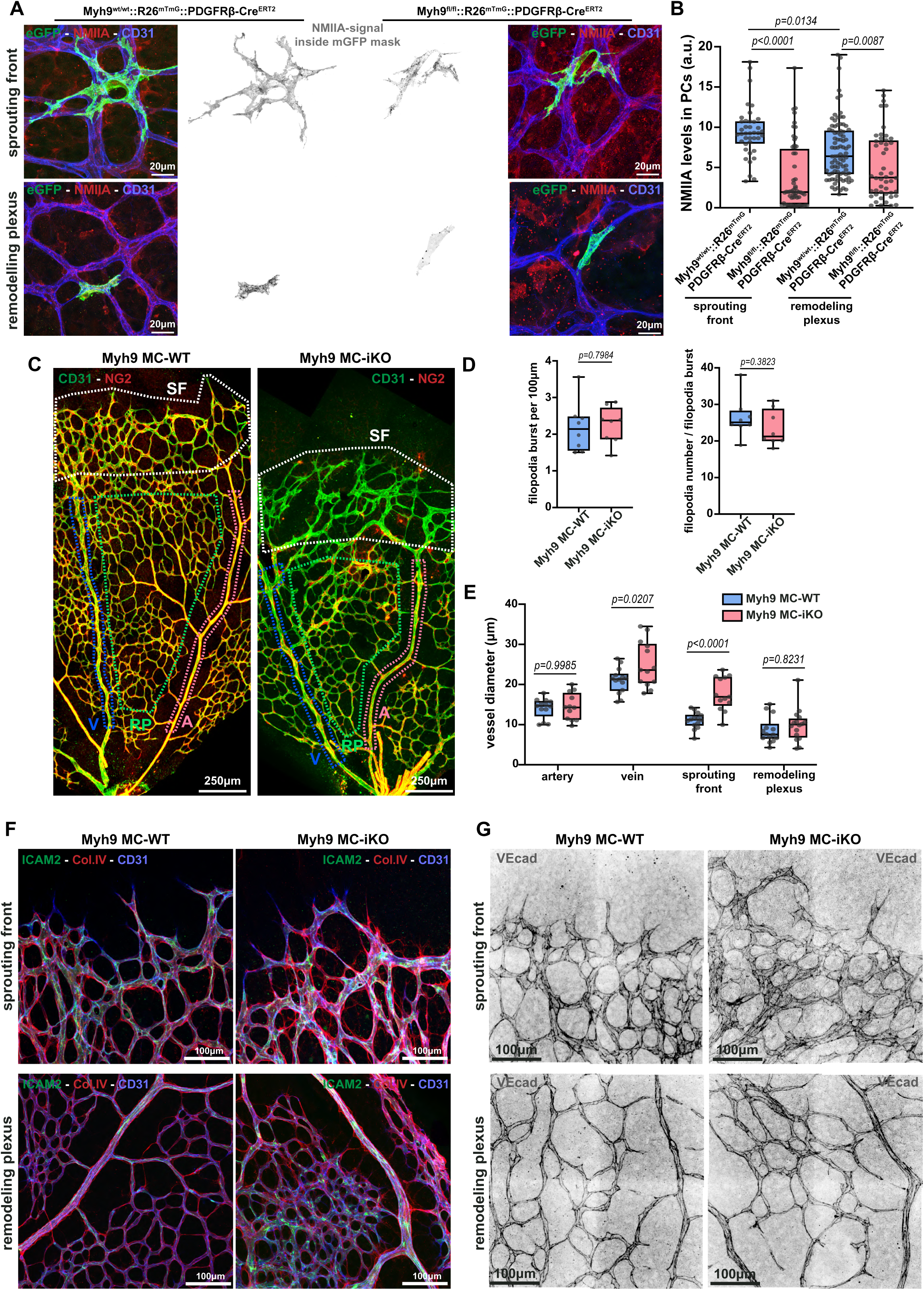
Structural vessel alterations after MC-specific Myh9 deletion. **A)** Representative images of SF and RP of P6 retinas from Myh9^wt/wt^::Pdgfrβ-Cre^ERT2^::R26^mTmG^ and Myh9^fl/fl^::Pdgfrβ-Cre^ERT2^::R26^mTmG^ mice. Single MCs (mGFP, green) are visualised within the capillary network (CD31, blue), together with non-muscle myosin IIA (NMIIA; red and grey). **B)** Quantification of NMIIA levels in MCs in the SF and RP of P6 retinas from Myh9^wt/wt^::Pdgfrβ-Cre^ERT2^::R26^mTmG^ and Myh9^fl/fl^::Pdgfrβ- Cre^ERT2^::R26^mTmG^ mice. Each dot represents one MC. SF: N=40 Myh9^wt/wt^::Pdgfrβ-Cre^ERT2^::R26^mTmG^, N=56 Myh9^fl/fl^::Pdgfrβ-Cre^ERT2^::R26^mTmG^; RP: N=89 Myh9^wt/wt^::Pdgfrβ-Cre^ERT2^::R26^mTmG^, N=48 Myh9^fl/fl^::Pdgfrβ-Cre^ERT2^::R26^mTmG^. **C)** Representative images of P6 retinas from *Myh9*^MC-WT^ and *Myh9*^MC-iKO^ mice. Retinas were stained for vessels (CD31, green) and PCs (NG2, red). Representative delineation of arteries (pink), veins (blue), the sprouting front (SF, white), and the remodelling plexus (RP, green) is shown in the figure. **D)** Quantification of filopodium bursts (left panel) and the number of filopodia (right panel) in the SF of P6 retinas from *Myh9*^MC-WT^ and *Myh9*^MC-iKO^ mice. Each dot represents one mouse retina. N=8 mice for both *Myh9*^MC-WT^ and *Myh9*^MC-iKO^. **E)** Quantification of vessel diameter in the specified vessel region in P6 retinas from *Myh9*^MC-WT^ and *Myh9*^MC-iKO^ mice. Each dot represents one vessel/capillary region. Artery: N=13 *Myh9*^MC-WT^, N=12 *Myh9*^MC-iKO^; Vein: N=14 *Myh9*^MC-WT^, N=13 *Myh9*^MC-iKO^; RP: N=14 *Myh9*^MC-WT^, N=16 *Myh9*^MC-iKO^; SF: N=14 *Myh9*^MC-WT^, N=13 *Myh9*^MC-iKO^. **F)** Representative images of collagen empty sleeves in the SF and RP of P6 retinas from *Myh9*^MC-WT^ and *Myh9*^MC-iKO^ mice, stained for vessels (CD31, blue), ICAM-2 (green), and collagen IV (ColIV, red). **G)** Representative images of the junctional marker VE-Cadherin (in grey) in the SF and RP of P6 retinas from Myh9 *Myh9*^MC-WT^ and *Myh9*^MC-iKO^ mice. All data are presented as box plots with whiskers, min to max, with median from at least 3 independent litters.

**Figure S6.**
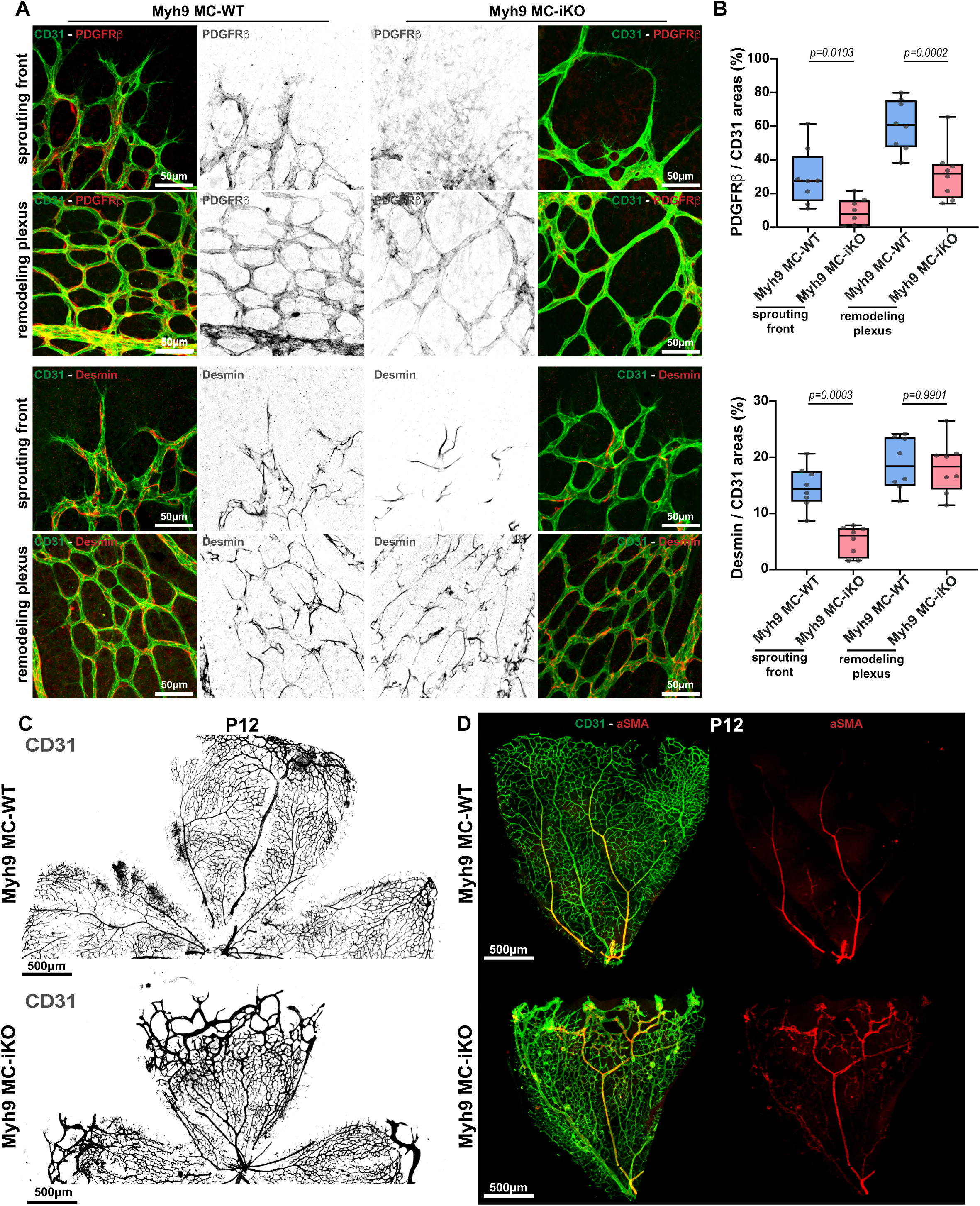
Altered PC coverage in Myh9 MC-iKO retinas. **A)** Representative images of PC markers PDGFRβ (upper panels) and desmin (bottom panels) in the SF and RP of P6 retinas from *Myh9*^MC-WT^ and *Myh9*^MC-iKO^ mice, stained for vessels (CD31, green) and PDGFRβ or desmin (red). **B)** Quantification of PC coverage (PDGFRβ+ or desmin+) in vessels (CD31+) in the SF and RP of P6 retinas from *Myh9*^MC- WT^ and *Myh9*^MC-iKO^ mice. Each dot represents one mouse retina. N=8 mice for both *Myh9*^MC-WT^ and *Myh9*^MC-iKO^. Data are presented as box plots with whiskers, min to max, with median from at least 3 independent litters. **C)** Representative images of blood vessels (CD31, grey) of P12 retinas from *Myh9*^MC-WT^ and *Myh9*^MC-iKO^ mice. **D)** Representative images of P12 retinas from Myh9^MC-WT^ and Myh9^MC-iKO^ mice stained for vessels (CD31, green) and αSMA (red), showing the expanded αSMA+ MC coverage associated with the arteriovenous shunt-like malformations of Myh9MC-iKO retinas.

**Figure S7.**
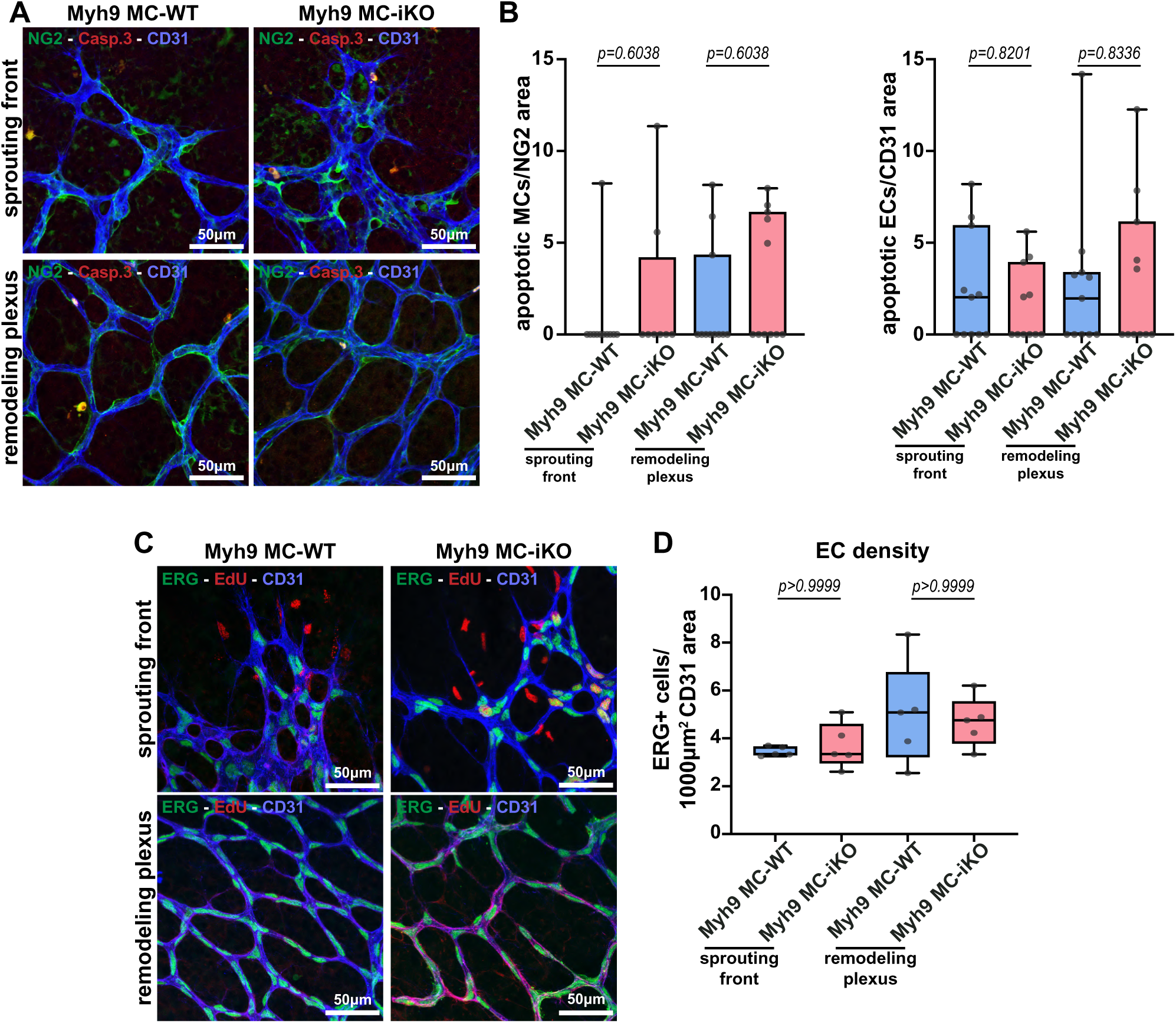
MC-specific Myh9 deletion does not affect cell apoptosis or EC density. **A)** Representative images of SF and RP of P6 retinas from *Myh9*^MC-WT^ and *Myh9*^MC-iKO^ stained for vessels (CD31, blue), PCs (NG2, green), and cleaved caspase-3 (Casp3, red). **B)** Quantification of apoptotic MCs (left panel) and ECs (right panel) at the SF and RP of P6 retinas from *Myh9*^MC-WT^ and *Myh9*^MC-iKO^ mice. Each dot represents one mouse retina. MCs, N=9 for both *Myh9*^MC-WT^ and *Myh9*^MC-iKO^. ECs, N=11 for *Myh9*^MC-WT^ and *Myh9*^MC-iKO^. **C)** Representative images of SF and RP of P6 retinas from *Myh9*^MC-WT^ and *Myh9*^MC-iKO^ mice stained for vessels (CD31, blue), ECs (ERG, green), and proliferating cells (EdU, red). **D)** Quantification of EC density in the SF and RP of P6 retinas from *Myh9*^MC-WT^ and *Myh9*^MC-iKO^ mice. Each dot represents one mouse retina. N=5 mice for both *Myh9*^MC-WT^ and *Myh9*^MC-iKO^. All data are presented as box plots with whiskers, min to max, with median from at least 3 independent litters.

**Figure S8.**
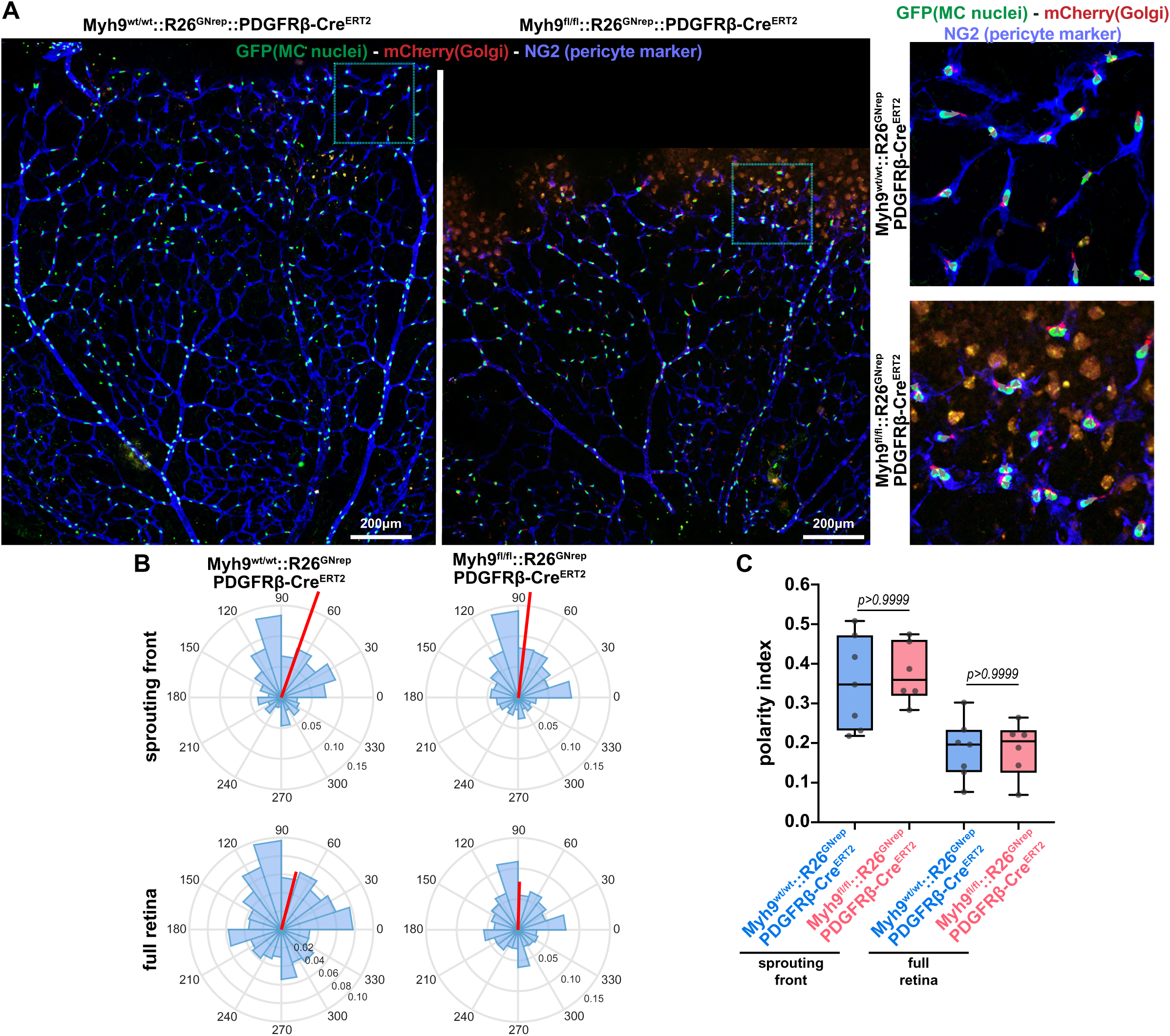
PC front-rear polarity is preserved upon MC-specific Myh9 deletion. **A)** Representative images of SF and RP of P6 retinas (left panels) from Myh9^wt/wt^::R26^GNrep^::Pdgfrβ-Cre^ERT2^ and Myh9^fl/fl^::R26^GNrep^::Pdgfrβ-Cre^ERT2^ mice stained for PCs (NG2, blue), and endogenously labelled Golgi (GTS-mCherry, red), and endogenously labelled nuclei (H2B-GFP, green). Right panels: higher-magnification images of the SF from corresponding inserts in left panel images. **B)** Rose plots showing MC polarity orientation at the SF or the entire retina from P6 Myh9^wt/wt^::R26^GNrep^::Pdgfrβ- Cre^ERT2^ and Myh9^fl/fl^::R26^GNrep^::Pdgfrβ-Cre^ERT2^ mice. **C)** Quantification of the MC front-rear polarity index at the SF and in the full retina from P6 Myh9^wt/wt^::R26^GNrep^::Pdgfrβ- Cre^ERT2^ and Myh9^fl/fl^::R26^GNrep^::Pdgfrβ-Cre^ERT2^ mice. Each dot represents one petal of a mouse retina. N=7 petals Myh9^wt/wt^::R26^GNrep^::Pdgfrβ-Cre^ERT2^, N=6 Myh9^fl/fl^::R26^GNrep^::Pdgfrβ-Cre^ERT2^. Data are presented as box plots with whiskers, min to max, with median from 3 independent litters.

**Figure S9.**
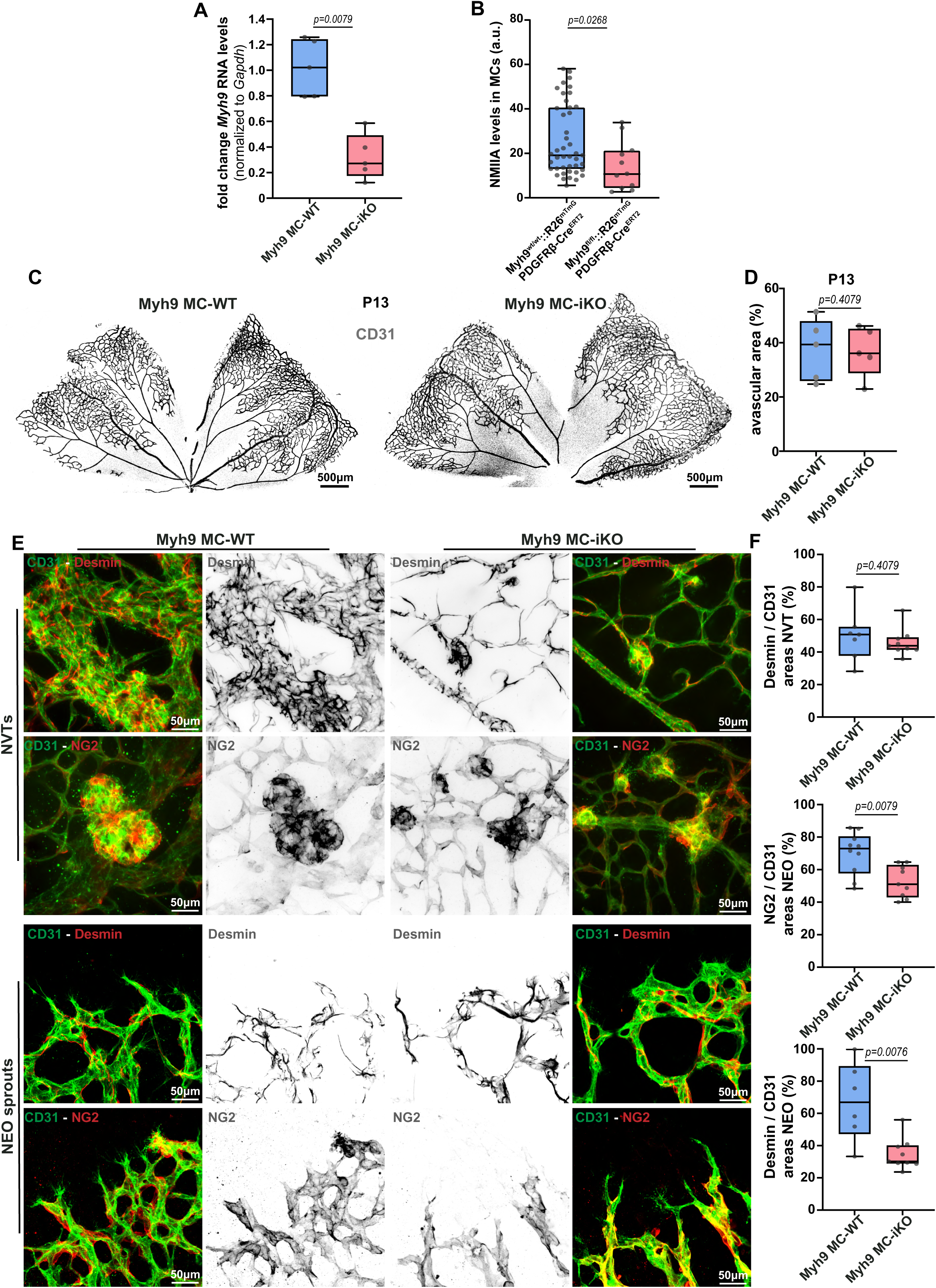
PC coverage in neovascular and neo-angiogenic regions after MC-specific deletion of Myh9. **^A)^** Myh9 mRNA expression levels in P16 OIR retinas from *Myh9*^MC-WT^ and *Myh9*^MC-iKO^ mice retinas. Each dot represents one animal. N=5 for both *Myh9*^MC-WT^ and *Myh9*^MC-iKO^. **B)** Quantification of NMIIA levels in MCs of P16 OIR retinas from Myh9^wt/wt^::Pdgfrβ- Cre^ERT2^::R26^mTmG^ and Myh9^fl/fl^::Pdgfrβ-Cre^ERT2^::R26^mTmG^ mice. Each dot represents one MC. N=43 Myh9^wt/wt^::Pdgfrβ-Cre^ERT2^::R26^mTmG^; N=11 Myh9^fl/fl^::Pdgfrβ-Cre^ERT2^::R26^mTmG^. **C)** Representative images of P13 (post-hyperoxia) retinas from Myh9^MC-WT^ and Myh9^MC- iKO^ mice stained for CD31 (grey). **D)** Quantification of the percentage of avascular area at P13 in Myh9^MC-WT^ and Myh9^MC-iKO^ retinas. Each dot represents one mouse retina. N=5 mice for *Myh9*^MC-WT^ and N=5 mice for *Myh9*^MC-iKO^. **E)** Representative images of NVTs (upper panels) or neovascular sprouts (NEO) (bottom panels) of P16 OIR retinas from *Myh9*^MC-WT^ (left panels) and *Myh9*^MC-iKO^ (right panels) mice. NVTs were stained for CD31 (green) and desmin (red and grey; top panel) or NG2 (red and grey; bottom panel). **F)** Quantification of the percentage of PC coverage (desmin) in individual NVTs, and of NG2 and desmin coverage in NEO (CD31+), of P16 OIR retinas from *Myh9*^MC-WT^ and *Myh9*^MC-iKO^ mice. For NVTs, each dot represents one NVT. N=7 *Myh9*^MC-WT^; N=9 *Myh9*^MC- iKO^ NVTs. For NEO, each dot represents one retina. N=10 *Myh9*^MC-WT^; N=9 *Myh9*^MC-iKO^ NVTs. All data are presented as box plots with whiskers, min to max, with median from at least 3 independent litters.

**Figure S10.**
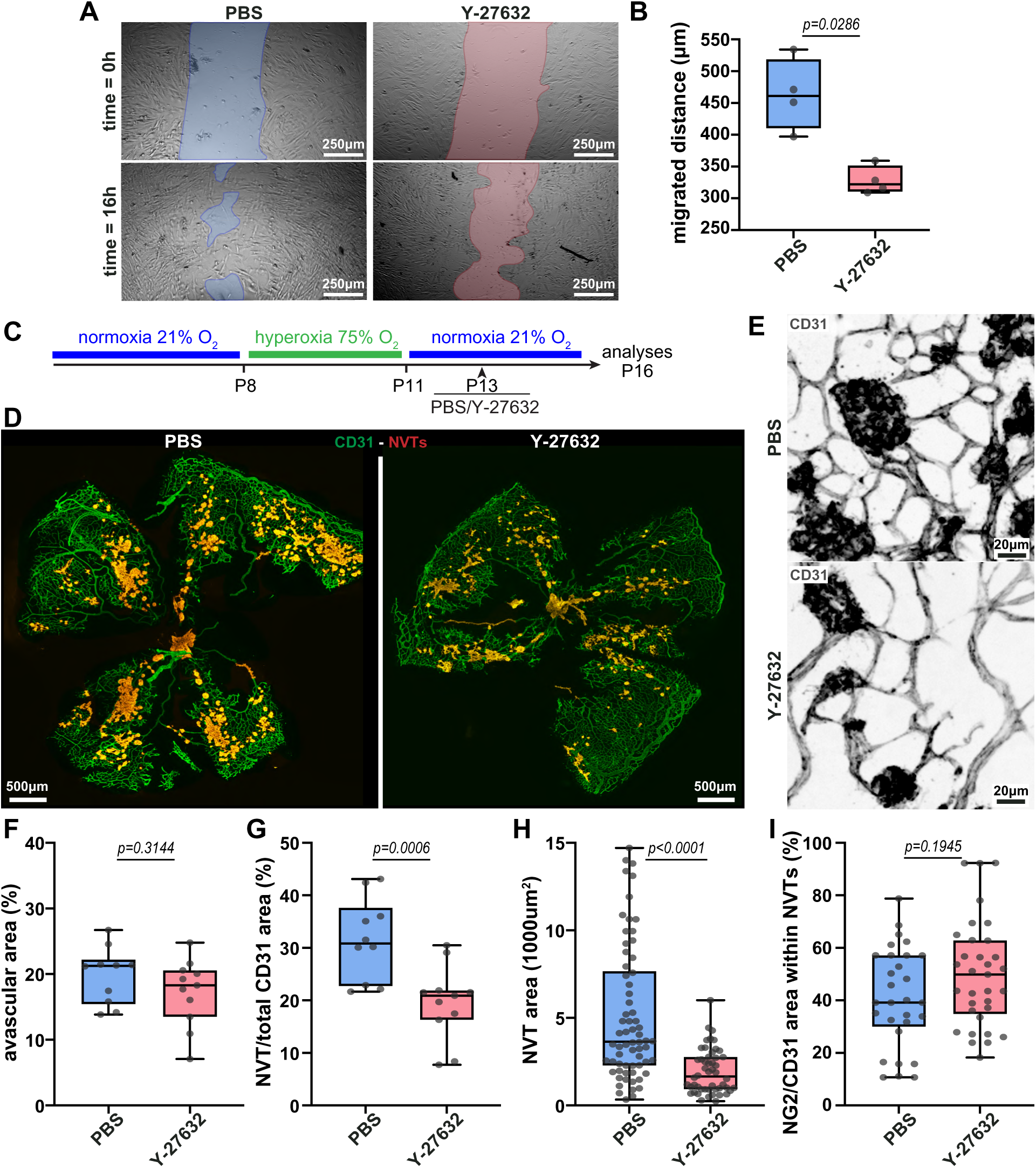
ROCK inhibition phenocopies MC-specific deletion of Myh9 in OIR. **A)** Representative images from a scratch-wound assay performed on hBPCs treated with PBS (control, left) or Y-27632 (ROCK inhibitor, right) at 0 and 16 hours. The wound area is shown at both time points. **B)** Quantification of the migrated distance of hBPCs treated with PBS or Y-27632. Each dot represents one well. N=4 PBS, N=4 Y-27632. **C)** Experimental timeline for ROCK inhibition during OIR. **D)** Representative images of P16 OIR retinas from PBS- or Y-27632-treated mice stained for CD31 (green). Neovascular tufts (NVTs) were manually delineated (red) and superimposed on CD31 staining**. E)** Higher-magnification images of NVTs stained for CD31 (grey) from PBS- or Y-27632- treated mice. **F)** Quantification of the percentage of avascular area in P16 OIR retinas from PBS- or Y-27632-treated mice. Each dot represents one mouse retina. N=10 PBS; N=10 Y-27632. **G)** Quantification of the total NVT burden per retina, calculated as the sum of all NVT areas divided by the whole-retina CD31+ area (NVT/CD31 areas), in P16 OIR retinas from PBS- or Y-27632-treated mice. Each dot represents one mouse retina. N=10 PBS; N=11 Y-27632. **H)** Quantification of the individual NVT area in P16 OIR retinas PBS- or Y-27632-treated mice. Each dot represents one NVT. N=66 PBS; N=50 Y-27632. **I)** Quantification of the percentage of PC coverage (NG2+) in individual NVTs (CD31+) of P16 OIR retinas from PBS- or Y-27632-treated mice. Each dot represents one NVT. N=30 PBS; N=33 Y-27632. All data are presented as box plots with whiskers, min to max, with median from at least 3 independent litters.

**Figure S11.**
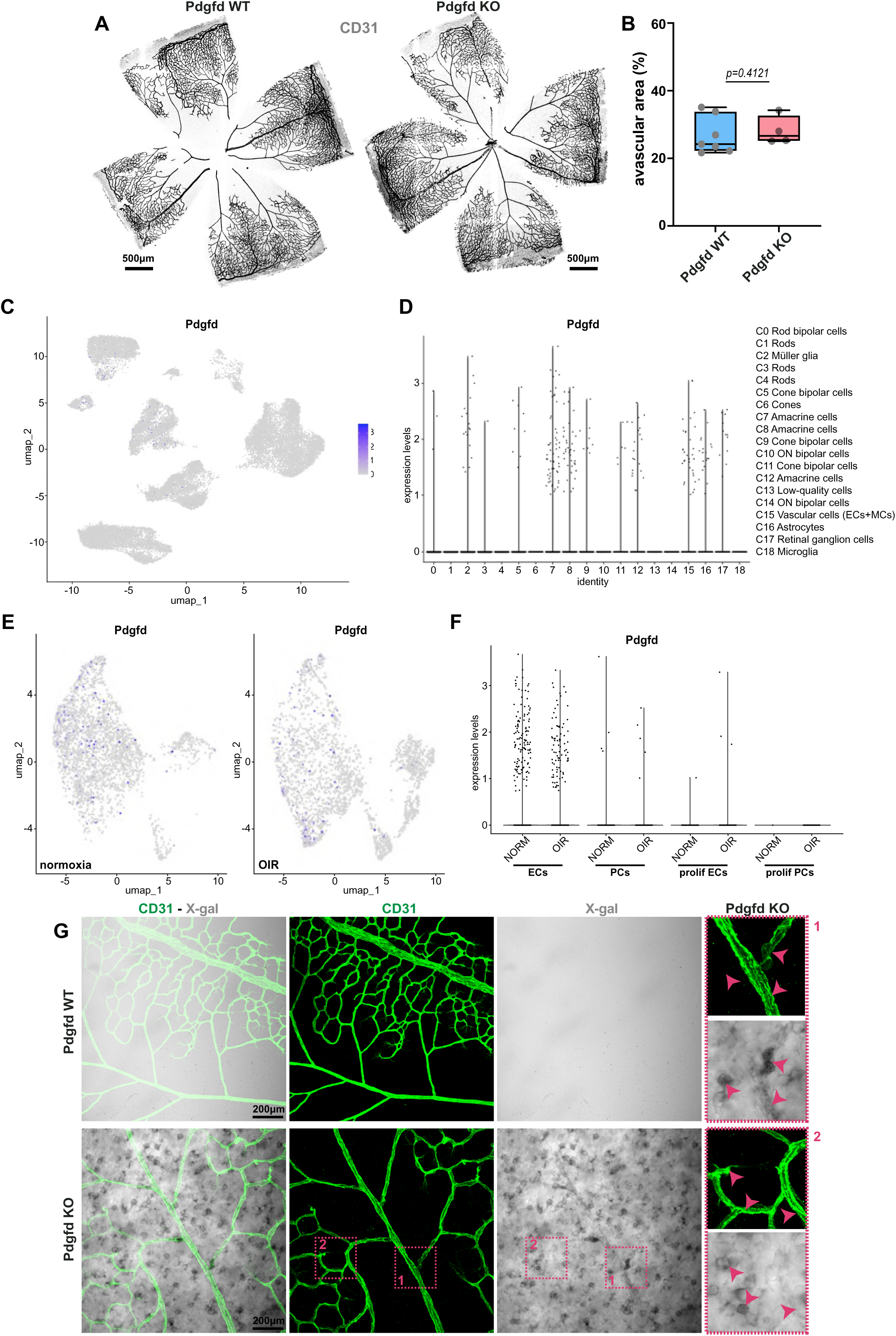
Pdgfd expression pattern in the mouse retina. **A)** Representative images of P13 (post-hyperoxia) retinas from Pdgfd^wt/wt^ (Pdgfd WT) and Pdgfd^-/-^ (Pdgfd KO) mice stained for CD31 (grey). **B)** Quantification of the percentage of avascular area at P13 in Pdgfd WT and Pdgfd KO retinas. Each dot represents one mouse retina. N=6 mice for Pdgfd WT and N=4 mice for Pdgfd KO. Data are presented as box plots with whiskers min to max, with median. **C)** Feature plot showing Pdgfd expression across all clusters of the retinal scRNAseq dataset. **D)** Violin plot showing Pdgfd expression level per designated cluster. **E)** Feature plots showing Pdgfd expression in the integrated EC/PC subset under normoxia (NORM) and OIR. **F)** Violin plots showing Pdgfd expression in ECs, PCs, proliferating ECs and proliferating PCs, split by condition (NORM, OIR). **G)** Representative images of retinas from Pdgfd WT and Pdgfd KO littermates stained with X-gal (brightfield, grey) and for CD31 (green). The Pdgfd-null allele carries a LacZ cassette, so that X-gal signal reports Pdgfd promoter activity and is present only in Pdgfd KO retinas. Insets 1 and 2 show higher magnifications of the boxed regions; arrowheads indicate X-gal+ cells, including ECs.

## SUPPLEMENTAL VIDEO LEGENDS

**Video S1.** Example of the three-dimensional segmentation and masking procedure used to quantify NMIIA in single cells. Three-dimensional rendering of a confocal stack of a P16 normoxic Cdh5-Cre^ERT2^::R26^mTmG^ retina showing, in sequence, the merged channels (CD31, blue; NMIIA, red; mGFP, green), the CD31 channel alone, the mGFP mask of the individual recombined cells, and the NMIIA signal contained within the mGFP mask. Related to Fig.S3B.

**Video S2.** Three-dimensional rendering of a single endothelial cell (mGFP; green) within vasculature (CD31+; blue) of a P16 normoxic Cdh5-Cre^ERT2^::R26^mTmG^ retina.

**Video S3.** Three-dimensional rendering of a single endothelial cell (mGFP; green) within a non-tuft capillary (CD31+; blue) of a D5 OIR Cdh5-Cre^ERT2^::R26^mTmG^ retina.

**Video S4.** Three-dimensional rendering of a single endothelial cell (mGFP; green) within a neovascular tuft (CD31+; blue) of a D5 OIR Cdh5-Cre^ERT2^::R26^mTmG^ retina.

**Video S5.** Three-dimensional rendering of a single pericyte (mGFP; green) within vasculature (CD31+; blue) of a D5 OIR PDGFRβ-Cre^ERT2^::R26^mTmG^ retina.

**Video S6.** Three-dimensional rendering of a single pericyte (mGFP; green) within a non-tuft capillary (CD31+; blue) of a D5 OIR PDGFRβ-Cre^ERT2^::R26^mTmG^ retina.

**Video S7.** Three-dimensional rendering of a single pericyte (mGFP; green) within a neovascular tuft (CD31+; blue) of a D5 OIR PDGFRβ-Cre^ERT2^::R26^mTmG^ retina.

## Notes

### Competing Interest Statement

The authors have declared no competing interest.

